# Remodelling of supernumerary leaflet primordia leads to bicuspid aortic valve (BAV) caused by loss of primary cilia

**DOI:** 10.1101/2025.02.25.636485

**Authors:** Ahlam Alqahtani, Lorraine Eley, Jake Newton, Kimberley Macdonald, Chloe Connolly, Cindy Rodrigues-Cleto, Kristyna Neffeova, Leonor Lopez, Javier Arias, Christopher J Derrick, Mashael Alaradi, Hana Kolesova, Bill Chaudhry, Deborah J. Henderson

## Abstract

**Aims:** Bicuspid aortic valve (BAV), where two valve leaflets are found instead of the usual three, affects 1-2% of the general population and is associated with significant morbidity and mortality. Despite its frequency, the majority of cases remain unexplained. This is, at least in part, because there are two types of valve leaflet primordia: endocardial cushions and intercalated valve swellings (ICVS). Moreover, multiple progenitors make distinct contribution to the formation of these primordia. Genomic studies in mouse and human have suggested a correlation between BAV and malfunctional primary cilia. However, the precise requirement for cilia during early embryonic valvulogenesis remains unknown.

**Methods and results:** Here, we disrupted primary cilia by deleting the ciliary gene Ift88 in the main progenitor cells forming the aortic valve using specific Cre drivers: Wnt1-Cre for neural crest cells, Isl1-Cre for second heart field cells (SHF); Tie2-Cre for endocardial-derived cells and Tnnt2-Cre for direct-differentiating SHF in the ICVS. Loss of Ift88, and thus primary cilia, from neural crest cells and endocardium did not impact aortic valve formation. However, primary cilia are essential in SHF cells for aortic valve leaflet formation, with over half of Ift88^f/f^;Isl1-Cre mutants presenting with BAV. As the valve leaflets are forming, 50% of the Ift88^f/f^;Isl1-Cre mutants have two small leaflets in the position of the usual posterior leaflet, meaning that at this stage the aortic valve is quadricuspid, which then remodels to BAV by E15.5. Mechanistic studies demonstrate premature differentiation of SHF cells as the ICVS form, leading to the formation of a broadened ICVS that forms two posterior leaflet precursors. This abnormality in the formation of the ICVS is associated with disruption of Notch-Jag1 signalling pathway, with Jag1^f/f^;Isl1-Cre mutants presenting with a similar phenotype.

**Conclusions:** These data show that primary cilia, via the Notch-Jag1 signalling pathway, regulate differentiation of SHF cells in the aortic valve primordia. Additionally, we identify a mechanistic link between the developmental basis of quadricuspid and bicuspid arterial valve leaflets.

**Translational Perspective:** Several genomic studies in human and mouse have suggested that disruption of cilia-related genes may be a significant cause of CHD. Although there is limited data from animal models to suggest a link between cilia and bicuspid aortic valve (BAV), the mechanisms underpinning BAV formation during early valvulogenesis have not been described. Here, we established a potential mechanism underpinning BAV formation, highlighting a role for primary cilia in a subset of valve interstitial cells (VIC) derived from second heart field progenitors. Loss of cilia altered VIC differentiation and valvulogenesis. This study confirms that disruption of cilial formation and/or function can lead to arterial valve defects and could pave the way to finding therapies for patient benefit.

## Introduction

Structural abnormalities of the heart are the commonest human congenital malformations. There is good evidence for an underlying genetic aetiology, but the mechanisms underlying the majority of these malformations remain undiscovered ^1^. It has been proposed that cilia, microtubular extensions found on almost all cells, at least in culture, may be important in vertebrate development. Motile cilia are responsible for generating fluid flows across the ciliated embryonic node which is fundamental to left-right specification ^2^. Disturbances of cilia function within the embryonic node are associated with reversal of organ positioning (situs inversus) and are likely to underly abnormal left-right patterning in some patients with complex congenital heart disease (heterotaxy). It has also been suggested that subtle disruption of left-right patterning may account for some apparently non-syndromic CHD ^3^. Distinctly, the non-motile primary cilium, found on almost all cells, acts as a hub for receptor-ligand signalling and force transduction ^4,5^. It has also been suggested that disruption of cilia-related genes may be a significant cause of CHD^6^.

Whilst genomic studies have revealed that variants in cilial genes are enriched in rare familial cases, this is not the case for isolated de novo cases of CHD ^7,8^. Patients with known mutations or syndromes affecting cilia usually present with kidney or eye disease but relatively rarely have CHD ^9^. More recently, family-based genetic studies have linked mutations in cilial genes to valve malformations such as bicuspid aortic valve (BAV) and mitral valve prolapse ^10,11^. This has been supported by similar findings in mouse studies ^6,12,13^.

*Ift88* (*intraflagellar transport 88*) is an intracellular transport protein required for ciliary biogenesis ^14^. In mice, complete knockout of *Ift88* is associated with a range of defects expected to occur following loss of cilia, including: laterality disturbances that impact the heart; neural tube abnormalities; polydactyly; and other retinal, kidney and brain defects ^15,16^. The hypomorphic Ift88 (*cobblestone*) mutant does not have situs inversus but exhibits a range of CHD including atrioventricular septal defect (AVSD), common arterial trunk (CAT) and hypoplasia of the ventricular myocardium ^17^. Tissue-specific knockout of Ift88 in the endocardium and its derivative cushion mesenchyme using Nfatc1-Cre resulted in BAV and thickened mitral valves ^11,13^ although the mechanisms underpinning these malformations during early valvulogenesis were not described. Thus, the published data suggests that primary cilia may be required in specific cell types within the developing outflow tract and valves of the heart.

Recent developmental studies have revealed many of the developmental processes that underlie valve formation. The leaflets found in the arterial valves have different embryological origins (Figure 1A). The left and right leaflets of both arterial valves (these are the coronary leaflets in the aortic valve) form from the distal part of the superior and inferior outflow tract cushions and are populated by invasion of neural crest cells (NCC) and endothelial cells undergoing endothelial to mesenchymal transformation (EndoMT; referred to here as endothelial derived cells; EDC). These latter cells are originally derived from second heart field (SHF) progenitors. The posterior (non-coronary leaflet) of the aortic valve, and the corresponding anterior leaflet of the pulmonary valve, form directly from SHF progenitors via intermediary structures known as intercalated valve swellings (ICVS) ^18,19^. We have previously reported that the ICVS differentiate directly from Isl1+ progenitor cells into Sox9 -expressing valve interstitial cells (VIC) via a Notch-Jag dependent mechanism ^18^. Thus, NCC, EDC (of SHF origin) and directly differentiating cells in the outflow wall (DD-SHF) all form VIC and are required for arterial valve morphogenesis. Our study, therefore, aimed to investigate the requirement for primary cilia in these different progenitor cells in the light of their various contributions to the formation of arterial valve leaflets.

**Figure 1:**
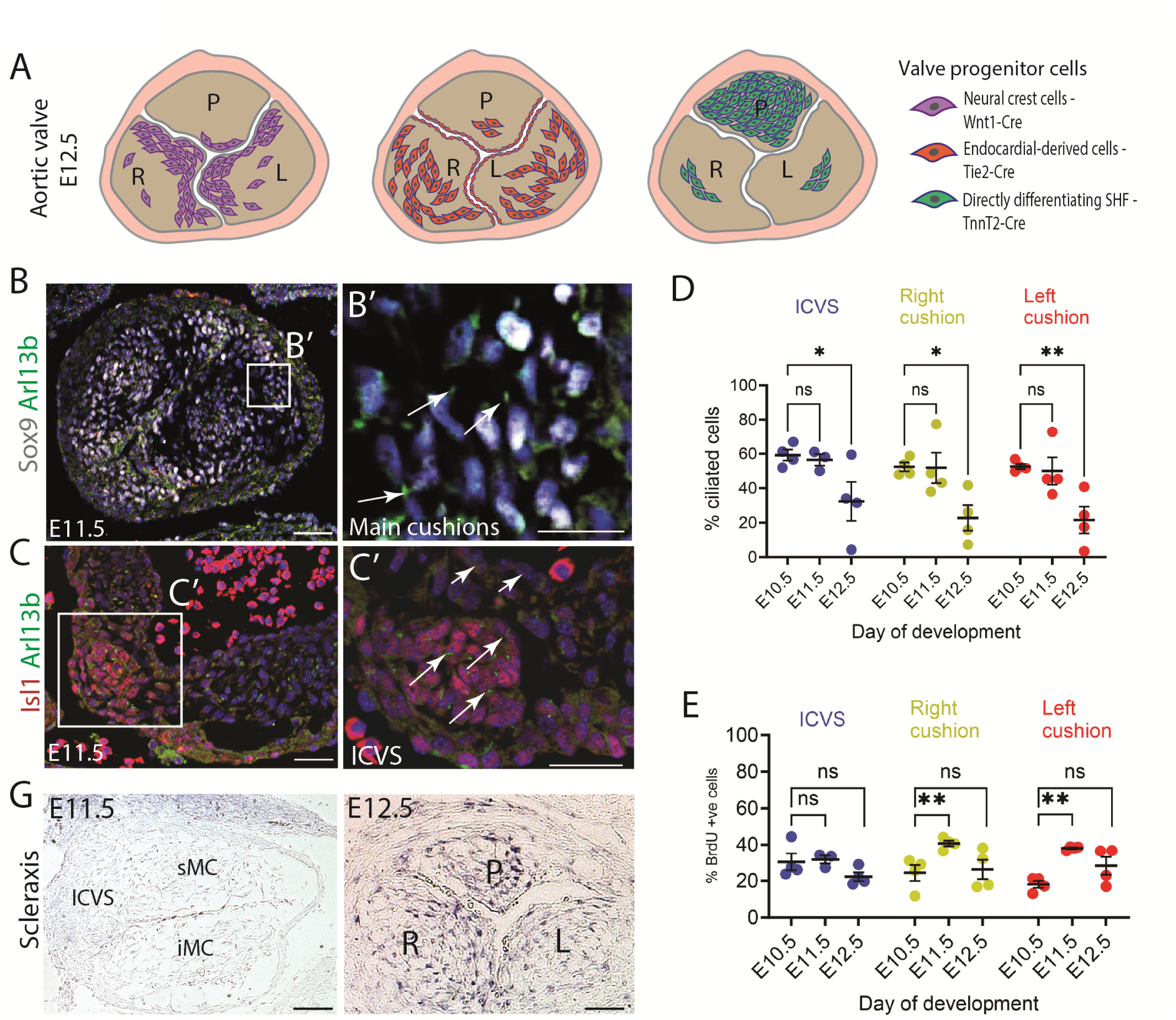
Dynamic presence of primary cilia in the main progenitor cells forming the arterial valves. **A)** Schematic view of the different cell linage contributions to the three leaflet primordia of the aortic valve at E12.5 [18, 19]. **B-B’)** Representative immunofluorescent images showing primary cilia marked by Arl13b (green; arrows in B’) found on the cells of the main outflow cushions (labelled by Sox9; white) at E11.5 (n=3). **C-C’)** Representative immunofluorescent images of primary cilia labelled by Arl13b (green; arrows in C’) found on the cells of the ICVS (labelled by Isl1; red) at E11.5 (n=3). In all cases, cell nuclei are labelled by DAPI (blue). **D)** The percentage of ciliated cells found in the ICVS, and the left and right main cushions/leaflets followed the same dynamic pattern from E10.5 to E12.5, with the majority of cells bearing a cilium up to E11.5, with a drop to 20-30% at E12.5 (n=4). Two-way ANOVA with Tukey test, Mean ± SD. ns: not significant, *p<.05, **p<.01. **E)** Quantification of proliferating cells labelled by BrdU showed no significant increase in proliferation at E12.5 that could account for the reduction in the percentage of ciliated cushion cells observed at E12.5 (n=4). Two-way ANOVA with Tukey test, Mean ± SD. ns: not significant, **p<.01. **F)** Absence of expression of Scleraxis in the main cushions (sMC and iMC) and ICVS of the outflow tract at E11.5 (n=3). **G)** Expression of Scleraxis in the aortic valve leaflets at E12.5, suggesting upregulation of VIC differentiation at this time point (n=3). ICVS: intercalated valve swellings, sMC: superior main cushion, iMC: inferior main cushion L: left leaflet, R: right leaflet, P: posterior leaflet. Scale bar = 100μm.

In this study, we show that loss of *Ift88* in SHF cells expressing Isl1-Cre leads to formation of quadricuspid valve leaflet primordia, that later remodel into a bicuspid valve. The loss of cilia leads to disruption of Notch signalling, suggesting that this pathway may be cilia-dependent for arterial valve development. Hence, this study uncovers a potential disease mechanism involved in the formation of BAV.

## Materials and Methods

### Mouse strains

*Ift88* flox (B6.129P2-Ift88tm1Bky/J;^20^) and *Jag1* flox (B6.129S-Jag1^tm2Grid^/SjJ;^21^) were obtained from Jackson Labs and genotyped as described previously. These were maintained on the C57Bl/6J background (Charles River Laboratories). *Wnt1-Cre* ^22^, *Tie2-Cre* ^23^, *Tnnt2-Cre*^24^, *Mef2c-AHF-Cre* ^25^ and *Isl1-Cre* ^26^ mice were inter-crossed with the *Ift88* flox mice to conditionally remove *Ift88* from the relevant lineage/cell type. *Jag1* flox mice were inter-crossed with *Isl1-Cre* mice to conditionally remove Jag1 from SHF cells. The reporter lines, *mTmG* ^27^ or the *ROSA-Stop-eYFP* ^28^, were also crossed into these lines to follow the relevant lineage/cell type. Timed mattings were carried out overnight with the Cre allele introduced by male mice; the presence of a copulation plug was designated embryonic day (E) 0.5. Littermate controls were used where possible. Each experiment was carried out, at minimum, in triplicate. All animals were euthanized by CO_2_ inhalation followed by a secondary physical method cervical dislocation. No anaesthesia was carried out on any animals in this study.

All mice were maintained according to standard laboratory conditions and all procedures were carried out in accordance with the local Animal Welfare and Ethical Review Body (AWERB), the UK Animals (Scientific Procedures) Act 1986, and Newcastle University (Project Licences 604533, P9E095FF4 and PP5418872) in line with Directive 2010/63/EU of the European Parliament. The Jag1;Isl1-Cre mouse study was approved by the local Animal Care and Use Committee of the First Faculty of Medicine, Charles University, in accordance with the local legislation and institutional requirements.

To calculate survival rates of different genotypes within litters, *Ift88^f/+^;Cre^+^* males were crossed with *Ift88^f/f^* females, thus 25% *Ift88^f/f^;Cre^+^*offspring were expected. In the case of *Wnt1-Cre*, Ift88^f/+^;Wnt1-Cre^+^ males were crossed with either *Ift88^f/+^* or *Ift88^f/f^*females, generating *Ift88^f/f^;Wnt1-Cre^+^* mutant offspring at an expected ratio of 1:8 (12.5%) or 1:4 (25%), respectively. Mutant survival rates were calculated based on these ratios, with a Chi square test used to determine divergence from the expected ratios.

### Histological analysis

Harvested tissues were rinsed in ice-cold phosphate-buffered saline (PBS) and fixed in 4% paraformaldehyde at 4⁰C, the duration of which depended on the age of the embryo, before paraffin embedding. Embryos were embedded in either a transverse, frontal or sagittal orientation. For each analysis carried out using embryos, a minimum of three embryos for each developmental stage and section orientation were used (biological replicates). The analyses were carried out as separate experiments and a minimum of 5-10 sections (technical replicates) were analysed for consistency. For basic histological analysis, paraffin-embedded embryos were sectioned and stained with haematoxylin and eosin (H&E; Thermoscientific, 6766010; Sigma, HHS-16) and Alcian blue (Therofisher scientific; 40046-0025) following standard protocols. In some cases, slides were processed for H&E staining following immunofluorescence. In these cases, the slides were washed in water for three minutes before staining with haematoxylin for 10 minutes and then eosin for 5 minutes. This was followed by dehydration through a series of increasing concentrations of ethanol, then Xylene, and finally mounting the slides with Histomount (National Diagnostics, HS-103).

### Immunofluorescence

Methodology has been published previously ^29^. Sections were cut from paraffin embedded embryos at 8 µm using a rotary microtome (Leica) in a transverse, frontal or sagittal orientation. Slides were de-waxed with xylene (National Diagnostics) and rehydrated through a series of ethanol washes. Following washes in PBS, antigen retrieval was performed by boiling slides in citrate buffer (0.01 mol/L) pH 6.0 for 5 minutes. Sections were blocked in 10% FCS at RT for 30 minutes and then incubated overnight at 4°C with the following primary antibodies diluted 1/100 in 2% FCS: acetylated-tubulin (Sigma-Aldrich; T5293), Ift88 (Merck, ABC932), Arl13b (Merck; SAB2700707), Isl1 (40.2D6, Developmental Studies Hybridoma Bank), GFP (abcam; ab13970, Origene, TP401), eNOS (Santa Cruz; sc-654), Sox9 (abcam; ab185230 & Bio-techne; AF3075), SM22 alpha (abcam; ab14106), cardiac Troponin I (HyTest; 4T21/2), Notch1ICD (Cell Signalling; 4147), Jagged1 (Cell Signalling; 2620), Notch2 (abcam; ab8296), Notch3 (abcam; ab23426), Lef1 (abcam; ab2230), Axin2 (abcam; ab32197), Caspase3 (abcam, 96615), Collagen I (abcam; ab21286), Versican (abcam; ab1032), Periostin (abcam; ab14041), BrdU (abcam; ab6326). After washing, sections were incubated at room temperature for one hour, with secondary antibodies (diluted 1/200 in 2% FCS) conjugated to either Alexa 488, Alexa 594, or Alexa 647 (Life Technologies). Fluorescent slides were then mounted with Vectashield Mounting medium with DAPI (Vector Labs). For fluorescent Notch1ICD staining, endogenous peroxidases were inhibited by treating slides with 3% H_2_O_2_ for 5 minutes, then the slides were treated as if for immunofluorescence. Immunoreactivity of fluorescent Notch1ICD was observed using TSA^TM^ Cyanine 3 System (Perkin Elmer NEL704A001KT). TBST was used for all washes and dilution of antibodies instead of PBS. Each experiment was carried out in triplicate using separate embryos (>3) and imaging multiple sections of the structure of interest where possible. Immunofluorescence images were collected using a Zeiss Axioimager Z1 fluorescence microscope equipped with Zeiss Apotome 2 (Zeiss, Germany). Acquired images were processed with AxioVision Rel 4.9 software.

### Proliferation assay

Pregnant female mice were injected with BrdU (Bromodeoxyuridine / 5-bromo-2’-deoxyuridine; 19-160, Merk), interaperitoneally (IP), at a concentration of 100 µg/g and were sacrificed after two hours of treatment. The embryos were collected and fixed in 4% paraformaldehyde at 4°C, then processed for immunofluorescence.

### Slide in situ hybridisation

Methodology was carried out as previously described ^18^. Digoxigenin (DIG)-labeled RNA antisense probes corresponding to the protein coding regions of Scleraxis (ENSG00000260428) were synthesized by T7 RNA polymerase (MAXIScript, Ambion) and in vitro transcription (pGEM_T easy, Promega), respectively. The slide in situ hybridization was performed using 10 mm paraffin-embedded sections of E11.5 and E12.5 mouse embryos. Slides were de-waxed with xylene and rehydrated with ethanol series washes. Slides were then fixed in 4% PFA for 20 minutes at room temperature prior to the treatment with proteinase K (10 mg/mL) for 10 minutes at 37°C. Post-fixation by 4% PFA was performed at room temperature for 5 minutes. Slides were then processed to the acetylation reaction for 10 minutes in solution containing 100 mM triethanolamine, 40 mM hydrochloric acid and 250mM acetic anhydride. After that, pre-hybridization was performed for 2 hours at 68°C using hybridization buffer (50% formamide, 5x SSC (pH6), 200 mg/mL T-RNA, 100 mg/mL heparin, 0.1% Tween-20); then hybridization with RNA probes was carried out for 16 hours. Following a series of washes, slides were blocked in 2% MABT (maleic acid buffer containing Tween 20) blocking solution (Roche) for 2 hours and incubated with anti-digoxigenin-AP, fab fragments (Roche) overnight at 4°C. Slides were then developed in NBT/BCIP solution for 18–36 hours at room temperature. The reaction terminated through incubation in 100% methanol for 5 minutes, followed by dehydration with washes in a serial ethanol dilution (20-30s per wash), incubation in xylene twice for 5 minutes, and finally mounting slides with Histomount (Agar scientific). A minimum of 3 embryos at each stage and genotype were analysed.

### RNAScope

RNAscope was performed using the RNAscope Multiplex Fluorescent Reagent Kit v2 from Bio-techne (cat# 323100) according to the manufacturer’s instructions. Briefly, embryos were fixed in 4% PFA at 4°C overnight, serially dehydrated using ethanol, and embedded in paraffin oriented for transverse/frontal/sagittal sections. The embryos were then sectioned at 8 µm using a Leica microtome and sections were mounted on positively charged slides (Marienfield). The slides were then heated to 60°C for 10 minutes, allowed to cool to room temperature, and dewaxed in xylene for 6 minutes before dipping in 100% ethanol for 4 minutes and then air drying. After that, slides were incubated with hydrogen peroxide for 10 minutes at room temperature and rinsed in water before soaking in boiling target retrieval solution for 20 minutes using a steamer. After rinsing the slides in 100% ethanol and allowing to air dry, protease plus solution was added to the sections and incubated for 30 minutes at 40°C. RNA probes of channel 2 (Ptch1, 402811-C2; Hey2, 404651-C2) were diluted in a 1 in 50 volume of channel 1 probe (Gli3, 445511; Hes1, 417701) and incubated at 40°C for two hours. All probes were from Bio-techne. Following several sequential amplification and blocking steps, the slides were incubated in TSA Vivid Fluorophore kit (7527/1 KIT, 7523/1 KIT, Bio-techne) at a dilution of 1 in 1000 using TSA diluent. Finally, slides were mounted using prolong Gold antifade mounting (Thermo-fisher Scientific) and stored at −20°C until imaged.

### 3D reconstruction from histological/Immunofluorescent sections

3D reconstructions of either H&E or immunofluorescence images were performed to elucidate and generate volumetric measurements of aortic valve leaflets at embryonic stages E11.5, E12.5, E13.5 and E15.5 using Amira software (Thermo-Fisher scientific, v). Briefly, 8 μm sections were used to generate images for 3D reconstructions. A minimum of 10 images throughout the entirety of the aortic valve was imported to Amira software for alignment and construction. Manual tracing of each individual leaflets using the Amira brush tool was followed by combining all traces to generate a 3D structure via the Amira generate Surface tool ^30^. A total of 16 mutants and 16 littermate controls were analysed for BAV from different embryonic stages. Quantification of leaflets at E12.5 was performed using the default measurements in Amira. Data was then copied for statistical analysis and creation of graphs in GraphPad Prism 10. Where leaflet volumes were calculated, the position of the commissure, even if displaced towards the lumen, was used as the boundary between primordia and where two leaflets were present in the position of the posterior leaflet, they were counted together.

### Quantification of the percentage of ciliated cells

Quantification of the percentage of cells carrying a cilium was performed on deconvoluted images (from Z-stack images) of the outflow tract at E10.5-E15.5 using the cell counter plugin in Fiji. Four representative images from three different embryos per developmental time point were used. Data was presented as the percentage of total cells bearing a cilium (percentage ciliated cells). Student’s two-tailed unpaired t-test was performed using GraphPad prism10.

### Quantification of transcript/protein expression

The expression of Gli3, Ptch1, Lef1, N1ICD, Jag1, Hes1, and Hey2 was quantified by counting the cells that showed the expression of these markers as dots within their cell nuclei/membrane, in a representative 250×250mm region of the ICVS of the aortic valve. The total nuclei number (marked with DAPI) within that region was also counted to represent the percentages of marker positive nuclei as a proportion of the total. Five sections from four controls and four *Ift88^f/f^;Isl1-Cre* mutant embryos at E10.5 and E11.5 were used to generate the data. Student’s (two-tailed unpaired) t-test was performed using Graphpad Prism 10 to compare between values.

### Cell counts and statistical analysis

The nuclei of cells were manually counted in the aortic valve leaflet primordia at E11.5 to E15.5, stained with H&E in which haematoxylin labelled cell nucleus for total cell counts, or Immunofluorescent in which DAPI used for total cell counts, or Isl1 and Sox9 for cushion mesenchymal cell counts, from each of the three discrete primordia. In each case, a minimum of six representative sections from six different embryos were counted per time-point. All sections were 8μm in thickness and covered distal, mid, and proximal regions of aortic valve primordia. The data was presented as total cell counts for each leaflet or for all three leaflets together. Student’s (two-tailed unpaired) t-test, a one-way ANOVA, or two-way ANOVA test were performed using Graphpad Prism 10.

## Results

### Primary cilia are present on the main progenitor cells forming the arterial valves

The presence of primary cilia on cells within the aortic valve primordia was confirmed using immunofluorescence with an Arl13b antibody at E10.5-E12.5 (Figure 1 B-D). The majority of mesenchymal cells within the main cushions of the outflow tract possessed cilia at E10.5 and E11.5, as did Isl1-expressing cells in the ICVS (Figure 1 B,C), but there was an approximately 50% reduction in the number of cells exhibiting cilia at E12.5, with a similar dynamic observed between the different primordia (Figure 1D). Cilia have been described in low shear stress areas on the endocardium of the chicken heart^31^, however, similar to Toomer et al., 2017 [13], we did not observe them in our study, possibly because the valve precursors are exposed to high levels of laminar shear stress. As there are both NCC and EDC in the main cushions, we used Wnt1-Cre and Tie2-Cre labelling to distinguish them and showed there were no differences in the percentage of ciliated cells between the two progenitors (Supplementary Figure 1 A-C). The deciliation event at E12.5 was not correlated with an increase in proliferation, as measured by incorporation of BrdU, indicating that it was not linked to cells dissembling their cilium as they entered mitosis (Figure 1E). However, it did correlate with upregulation of the expression of *Scleraxis* in the cushions at E12.5 (Figure 1 F,G), suggesting it may be linked to differentiation of progenitors to VIC ^32^. Supporting this explanation, similar correlations between deciliation and differentiation were seen for myocardium and smooth muscle cell differentiation in the developing outflow tract (Supplementary Figure 1 D,E).

### Deletion of Ift88 in EDC and DD-SHF, but not NCC, results in cardiac abnormalities

We next investigated the requirement for primary cilia in the developing outflow tract by conditional deletion of Ift88, as it is required for ciliogenesis and is present in primary cilia of outflow cushion cells (Supplementary Figure 1F). In Ift88^f/f^;Wnt1-Cre (conditional knockout in neural crest cell), all fetuses examined (8/8) had craniofacial defects as previously described ^33^ (Figure 2 A,B). However, despite a reduction in the number of ciliated NCC within the pharyngeal arches of Ift88^f/f^;Wnt1-Cre mutants at E11.5 (Supplementary Figure 2A) no cardiovascular abnormalities were observed (Figure 2C). As this was a surprising finding, we checked whether NCC in the outflow tract cushions normally express Ift88, confirming that they do (Supplementary Figure 2C). Thus, the absence of expression of Ift88 by cardiac NCC was not an explanation for the lack of a phenotype. Furthermore, the numbers of Ift88^f^;Wnt1-Cre embryos collected at E15.5 did not suggest there was intrauterine loss (Supplementary Figure 2B). Thus, the loss of Ift88 in NCC disrupted cilia formation and craniofacial development but had no effect on outflow tract development.

**Figure 2:**
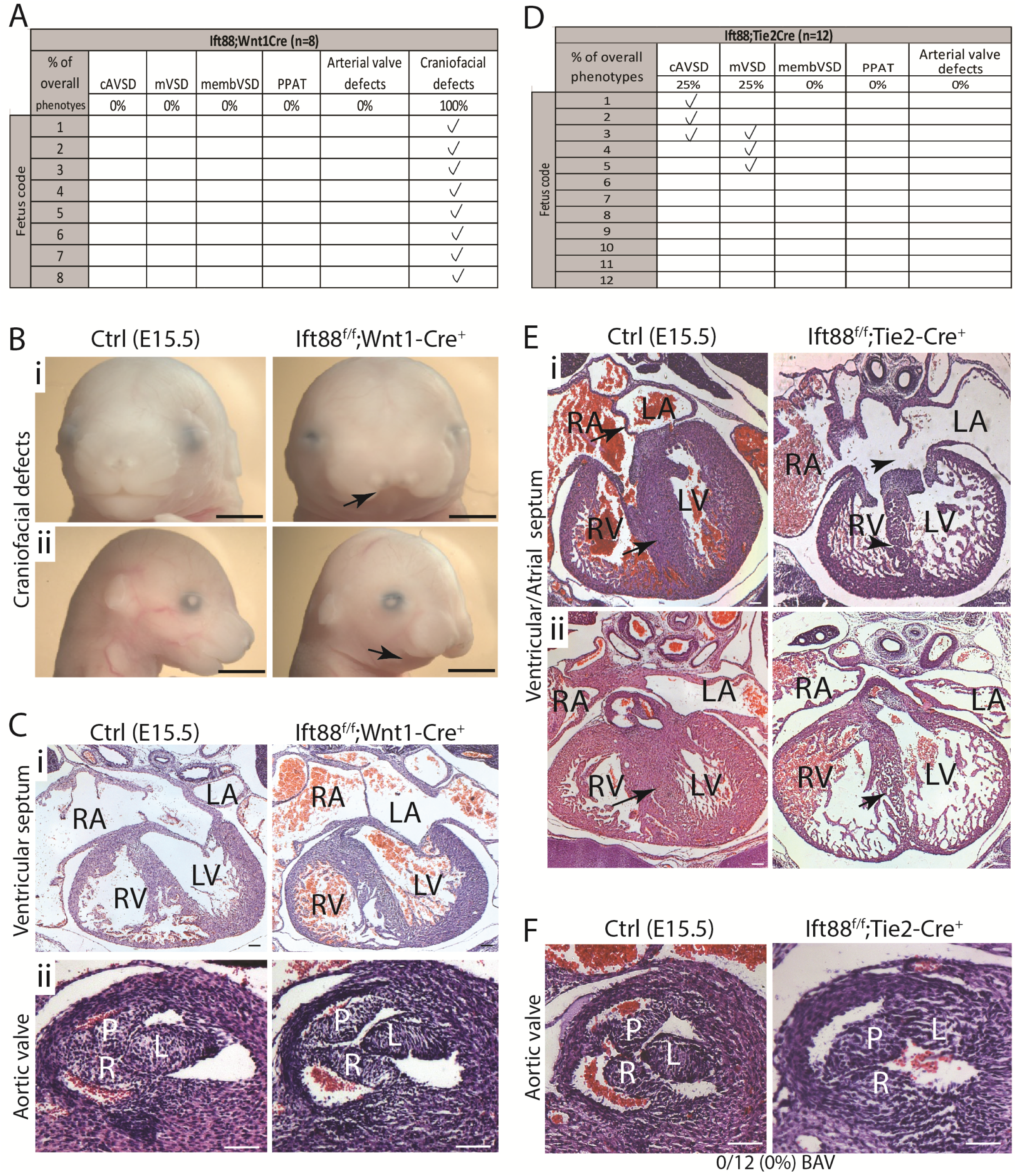
Primary cilia are not required in NCC or EDC-derived cells for aortic valve development. **A)** Table showing the different phenotypes observed within Ift88^ff^;Wnt1-Cre mutant fetuses at E15.5. **B)** Craniofacial defects including frontonasal dysplasia (arrow in i) and hypoplastic mandible (arrow in ii) were observed in Ift88^f/f^;Wnt1-Cre mutants (n=8/8) at E15.5. **C)** Histological analysis of the hearts of E15.5 Ift88^f/f^;Wnt1-Cre fetuses showed no observable differences in ventricular septation (i) or in the aortic valve (ii) between Ift88^f/f^;Wnt1-Cre mutants and control littermates. n=8 for each genotype. **D)** Table showing cardiac malformations observed in Ift88^ff^;Tie2-Cre mutant fetuses at E15.5. **E)** Histological analysis of Ift88^f/f^;Tie2-Cre mutants (n=12) showing complete AVSD (arrowheads in i, n=3/12) and muscular VSD (arrowhead in ii, n=3/12). Arrows show the normal structures in control littermates. **F)** The Ift88^ff^;Tie2-Cre aortic valve was comparable to control littermates in all fetuses examined (n=12 for each genotype). cAVSD, complete atrioventricular septal defect; mVSD, muscular ventricular septal defects; membVSD, membranous ventricular septal defect; PPAT, parallel proximal arterial trunk; P, posterior leaflet; R, right leaflet; L, left leaflet; LV, left ventricle; RV, right ventricle; LA, left atrium; RA, right atrium. Scale bar= 100mm (C,E,F) or 1mm (B).

We next investigated whether loss of Ift88 in the endocardial lineage, using Tie2-Cre, would disrupt cardiac, and particularly arterial valve, morphogenesis. In contrast to the Wnt1-Cre experiments, 42% of Ift88^f/f^;Tie2-Cre mutants (5/12) exhibited cardiac defects (Figure 2 D,E). Three mutants had complete atrioventricular septal defects (AVSD). Moreover, muscular ventricular septal defects, in the context of a thinned ventricular myocardium, were seen in 3/12 mutants (one of which also had AVSD). At E11.5, the atrioventricular cushions were underdeveloped and remained unfused in 2/6 cases, although the dorsal mesenchymal protrusion and primary atrial septum appeared well formed and fused with the superior atrioventricular cushion in all cases (Supplementary Figure 2D). Notably, there were no obvious abnormalities of the arterial valves in any of the twelve Ift88^f/f^;Tie2-Cre fetuses examined at E15.5 and the outflow cushions in mutants were comparable with controls at E11.5 (Figure 2F, Supplementary Figure 2D). There was no evidence of that Ift88^f/f^;Tie2-Cre mutants were dying before E15.5 (Supplementary Figure 2B).

In the Ift88^f/f^;Tnnt2-Cre conditional knockout, which targets DD-SHF cells found in the ICVS, eight Ift88^f/f^;Tnnt2-Cre mutant fetuses were analysed for cardiac defects (Figure 3A). Two had muscular VSDs, with one also having a membranous VSD, but there were no AVSD. An additional fetus had parallel arterial trunks but no VSD (Figure 3B). The aortic valve leaflets appeared thickened in 2/8 of Ift88^f/f^;Tnnt-Cre mutants (Figure 3C) and in one of these cases, the right leaflet of the aortic valve had a cleft distally (confirmed by three-dimensional (3D) reconstruction; Figure 3C). The pulmonary valve in one of the Ift88^f/f^;Tnnt2-Cre mutants also had thickening of all three leaflets; the remaining 7/8 mutants had a comparable pulmonary valve to their control littermates (Supplementary Figure 2E).

**Figure 3:**
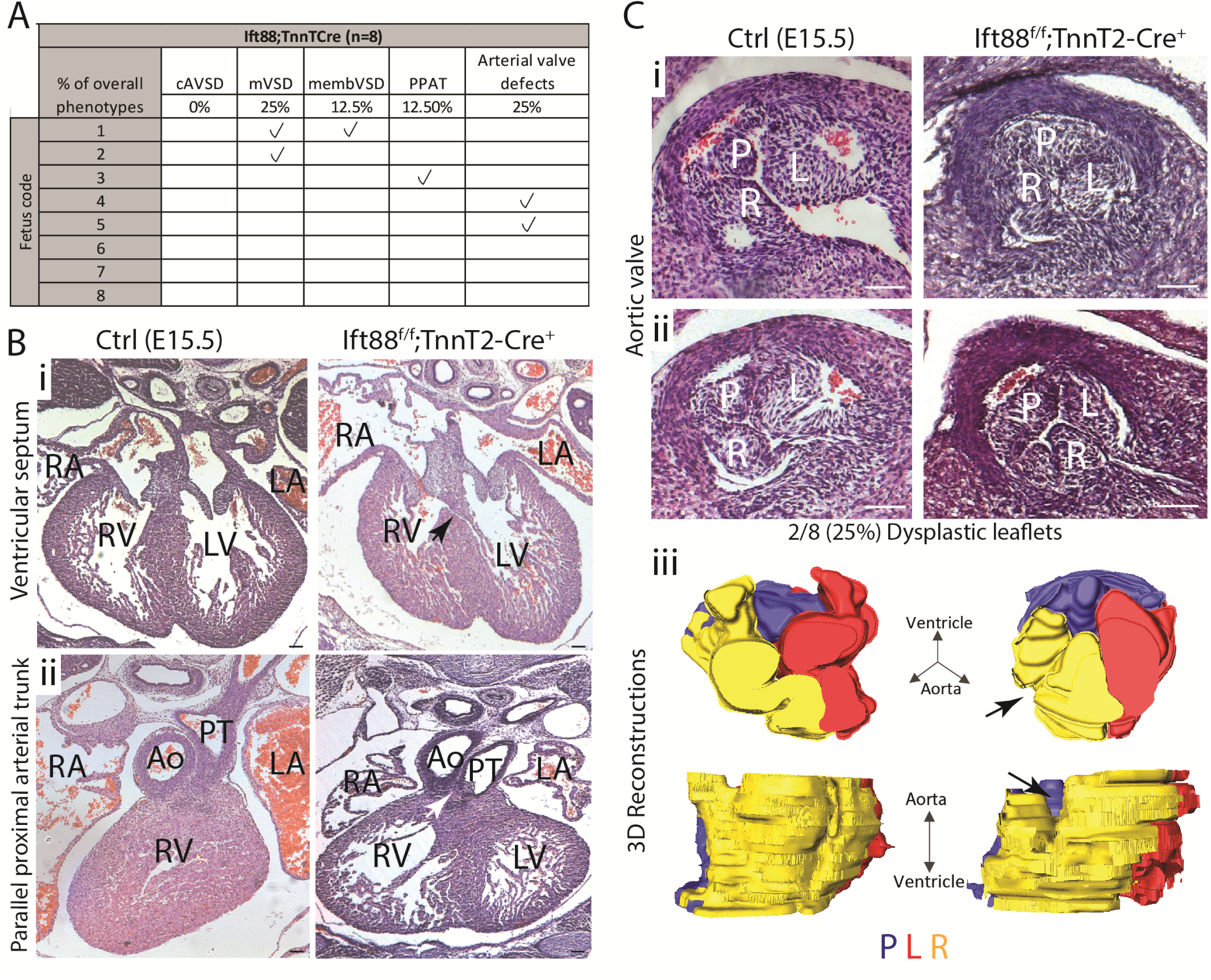
Primary cilia are required in the directly differentiating SHF cells for the aortic valve development. **A)** Table showing cardiac malformations observed in Ift88^ff^;Tnnt2-Cre mutant fetuses at E15.5. **B)** Morphological analysis of Ift88^f/f^;Tnnt2-Cre^+^ mutants and control littermates at E15.5 revealed several heart defects including muscular VSD (arrowhead in i) in 2/8 cases, and parallel proximal arterial trunks (white arrowhead in ii) in 1/8 mutants. **C)** Dysplastic aortic valve leaflets were observed in 2/8 mutants examined (i). In one of the E15.5 Ift88^f/f^;Tnnt2-Cre mutants, an apparent cleft was observed in the right leaflet (ii). Three-dimensional reconstruction confirmed the cleft in the distal region of the right leaflet (arrows in iii). cAVSD, complete atrioventricular septal defect; mVSD, muscular ventricular septal defects; membVSD, membranous ventricular septal defect; PPAT, parallel proximal arterial trunk; P, posterior leaflet; R, right leaflet; L, left leaflet; LV, Left ventricle; RV, right ventricle; LA, left atrium; RA, right atrium; Ao, Aorta; PT, pulmonary trunk. Scale bar= 100μm.

Together these data show that there were no arterial valve malformations when Ift88 was deleted in NCC or in the EDC, but there were dysplastic arterial valve leaflets in some cases when Ift88 was deleted in the Tnnt2-Cre lineage. This suggests that cilia may be required for the development of the arterial valve leaflets, but not primarily in the NCC or EDC lineages.

### Deletion of Ift88 in the Isl1Cre-targetted SHF leads to BAV

We next wanted to establish if earlier deletion of Ift88 (and thus cilia), in the undifferentiated SHF would result in arterial valve defects. We first crossed Mef2c-AHF-Cre mice with the Ift88^f^ line. No arterial valve abnormalities observed, although membranous VSD were seen in 2/8 mutant fetuses (Supplementary Figure 3A), as previously described in ^12^. There was no evidence of intrauterine loss of Ift88^f/f^;Mef2c-AHF-Cre mutants (Supplementary Figure 3B).

To examine an earlier requirement for Ift88 in the SHF, we turned to the Isl1-Cre line as this SHF lineage driver has a broader expression domain and is expressed earlier than Mef2c-AHF-Cre ^34^. Crosses between Ift88^f^ and Isl1Cre were carried out as for the other Cre lines. Cardiac abnormalities were present in 11 of 14 Ift88^f/f^;Isl1-Cre fetuses examined histologically at E15.5 (Figure 4A). Of these 11 mutants, one had double outlet right ventricle, one had muscular VSD and seven had membranous VSD (Figure 4 A,B); there were no cases of AVSD. Distinct from all other conditional knockouts of *Ift88*, BAV was apparent in 8/14 Ift88^f/f^;Isl1Cre mutant fetuses examined (Figure 4D, compare with 4C). Closer examination (that included 3D reconstruction) revealed that in all 14 Ift88^f/f^;Isl1-Cre mutants, there was a degree of disruption to the aortic valve leaflets, primarily affecting the posterior DD-SHF-derived leaflet. In five cases, the posterior leaflet was completely absent with a large commissure in the position where the missing leaflet would normally be located (Figure 4D). In three cases, there was an abnormally small posterior leaflet found only in the proximal part of the valve, with no associated sinus (Figure 4E). In the remaining six, whilst these were tri-leaflet aortic valves, the right coronary leaflet was thickened, and the posterior leaflet appeared to be mal-positioned and shorter (not extend the full length of arterial root) compared to controls (Figure 4F). Notably, Arl13b-expressing cilia were completely lost from Isl1+ SHF cells in the developing ICVS, although many cells (presumably NCC) still carried cilia in the main cushions (Figure 4 G). Quantification of the total cell counts revealed that although there were various degrees of aortic valve malformation, the total number of cells was comparable between controls and mutants (Figure 4H); suggesting malformation might not be caused by changes to overall cell numbers but rather in the allocation of cells within the leaflets. Moreover, there were no reproducible differences in the expression of extracellular matrix molecules such as Collagen I, Versican or Periostin at E18.5 or later at P1 (Supplementary Figure 4A), as has been reported for Ift88;Nfatc1-Cre mutants^13^.

**Figure 4:**
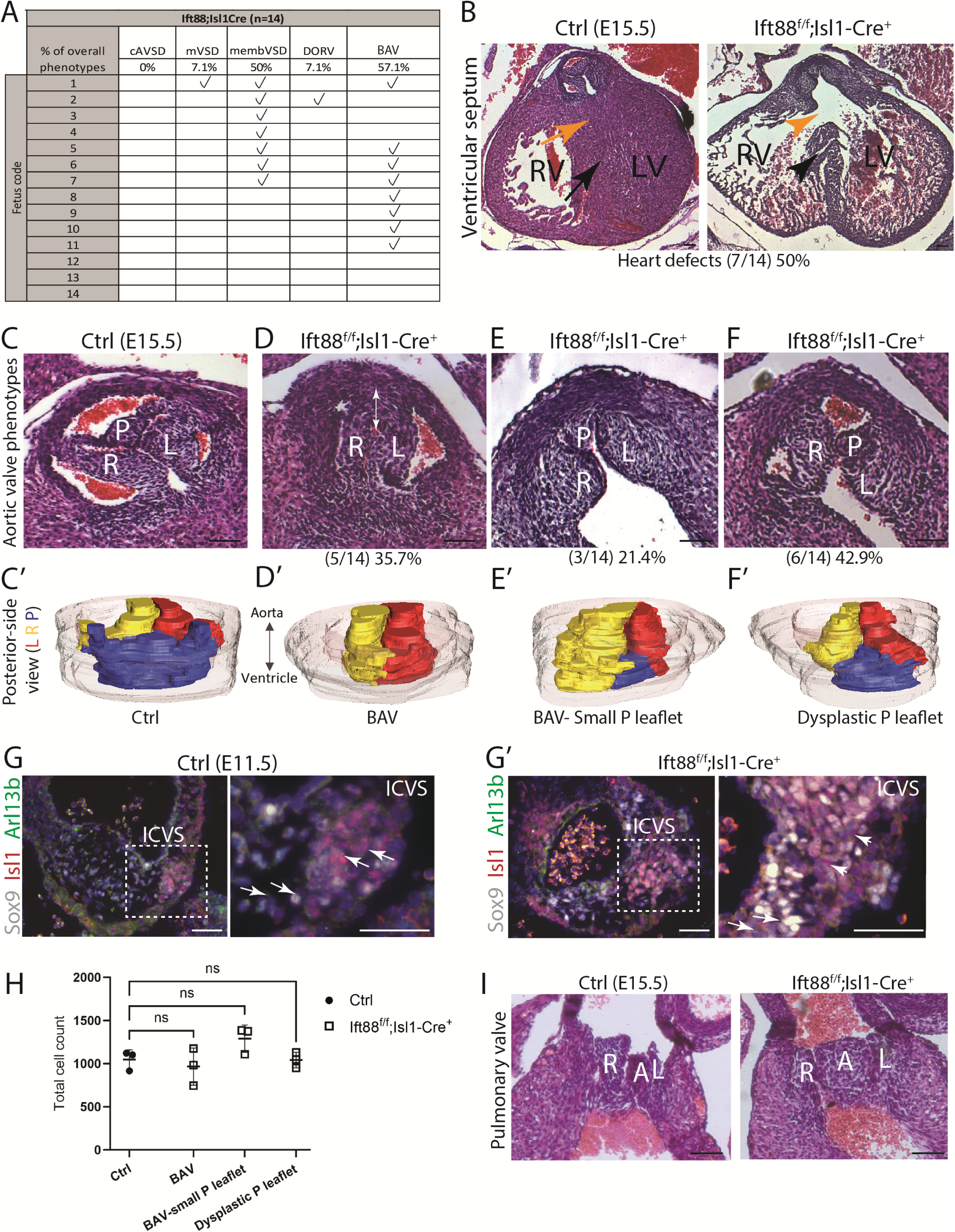
Loss of primary cilia in the SHF leads to abnormal aortic valve development. **A)** Table showing cardiac malformations observed in Ift88^ff^;Isl1-Cre mutant fetuses at E15.5. **B)** Histological analysis of Ift88^f/f^;Isl1-Cre at E15.5 (n=14) revealed several congenital heart defects including membranous VSD (7/14; yellow arrow and arrowhead) and muscular VSD (1/14; black arrow). **C-F)** BAV was observed in 8/14 of the Ift88^f/f^;Isl1-Cre fetuses at E15.5. Closer analysis of Ift88^f/f^;Isl1-Cre^+^ mutants, using histological sections and 3D reconstruction, revealed a spectrum of aortic valve malformations mainly affecting the posterior leaflet. In 5/8 mutants with BAV, the posterior leaflet was completely absent, with a large commissure found in the position of the missing leaflet (D-D’; white double headed arrow). In 3/8 of these mutants, the posterior leaflet only appeared in the proximal region of the valve, as a small leaflet with no associated sinus (E-E’). In the remaining 6/14 mutants with three leaflets, the posterior leaflet appeared dysplastic and the right leaflet relatively thickened. 3D reconstructions of the aortic valve showed that the posterior leaflet was shorter than the other two leaflets (compare F’ with C’). **G,G’)** Arl13b-expressing cilia are found on Isl1-expressing and Sox9-expressing cells in the ICVS in control embryos (G), whereas there are few if any cilia on the more abundant Sox9-expressing cells in the Ift88^f/f^;Isl1-Cre at comparable stages (G’). **H)** Quantitation of total cell counts in the aortic valve leaflets across the various abnormalities identified no significant changes in the overall cell number at E15.5, n=3 per group. One-way ANOVA with Tukey test, Mean ± SD. ns: not significant. **I)** Histological analysis of the pulmonary valve in Ift88^f/f^;Isl1-Cre^+^ mutants showed dysplastic anterior leaflet in 3/14 mutants. cAVSD, complete atrioventricular septal defect; mVSD, muscular ventricular septal defects; membVSD, membranous ventricular septal defect; DORV, double outlet right ventricle; L, left leaflet; P, posterior leaflet; R, right leaflet; A, anterior leaflet; LV, Left ventricle; RV, right ventricle. Scale bar=100μm.

No cases of bicuspid pulmonary valve were observed but dysplastic pulmonary anterior leaflets were present in 3/14 cases examined (Figure 4I). As for the other lines, there was no evidence of death prior to E15.5 (Supplementary Figure 3B).

To explore why the deletion of Ift88 using Isl1-Cre produced arterial valve abnormalities whilst Mef2c-AHF-Cre did not, we examined the expression patterns of two Cre drivers within the developing outflow tract and arterial valves. Between E10.5 to E11.5, when the ICVS were first apparent, Isl1-Cre labelled all the cells in the ICVS as well as the myocardial and endocardial cells that overlay the ICVS (Supplementary Figure 3 C-D). In contrast, while Mef2c-AHF-Cre labelled this myocardium and the endocardium, it did not label some cells within the ICVS (white and green arrowheads in Supplementary Figure 3 C,D). By E12.5, there was a subpopulation of cells in the endocardium and the underlying mesenchyme of the distal region of the posterior leaflet that was labelled by Isl1-Cre, but not Mef2c-AHF-Cre (Supplementary Figure 3E). Therefore, there are differences in the expression domains of Isl1-Cre and Mef2c-AHF-Cre that could potentially account for the differences observed when the different SHF-specific Cres are used to delete Ift88. Interestingly, the same region was labelled by both Cres in the pulmonary valve (Supplementary Figure 3F) suggesting that this cannot explain the milder defects observed in the pulmonary valve in Ift88^f/f^;Isl1-Cre fetuses. Therefore, although the differences in the expression patterns between Isl1-Cre and Mef2c-AHF-Cre could explain the differences in phenotypes, it could not account for the observed milder phenotypes in the pulmonary valves of Ift88^f/f^;Isl1-Cre.

Two leaflets are found in the position of the posterior leaflet in Ift88^f/f^;Isl1-Cre mutants Having taken a systematic approach, knocking out Ift88 in NCC and EndMT-derived cells that together contribute to the mesenchyme of the main cushions, we showed that there was no aortic valve phenotype in either case and thus loss of cilia from the main cushions did not lead to BAV. Thus, although deletion of Ift88 using Isl1Cre would affect EDC [24] and potentially some NCC^35^, the valve anomalies were very unlikely to be due to the role of Ift88 in the main cushions. Thus, by a process of genetic subtraction, we homed in on the other Isl1-Cre-derived arterial valve precursors – the ICVS. We started by observing the morphological structure of the aortic ICVS at E11.5, as it is the precursor of the posterior leaflet that was mostly affected in Ift88^f/f^;Isl1-Cre mutants at E15.5.

Histological sections of the E11.5 outflow tract identified that the posterior ICVS was present in all the mutants (n=14/14), but in the majority of cases appeared to be split into two smaller structures (n=12/14; Figure 5A). Cell counting for each of the primordia (cushions and ICVS) revealed no significant differences between controls and Ift88^f/f^;Isl1-Cre, suggesting that the defects were due to abnormal arrangement of cells, rather than an increase in cell number (Figure 5B). As at E11.5 the ICVS are composed of Isl1+ SHF progenitors that are differentiating to Sox9-expressing VIC ^18^, we immunolabeled the cells in the ICVS with these markers. The immunostaining of the aortic ICVS with Isl1 (red) and Sox9 (green) supported the appearances of two foci in the mutants compared to the expected single focus in the control littermates. Moreover, whereas Sox9+ cells were found only centrally (at the vertex of the oval-shaped, frontally-cut outflow tract) in the wild type embryos, they were found more laterally in the mutants resulting in the appearance of a broadened IVCS region (Figure 5C). Whilst the Isl1 and Sox9-expressing cells in the control ICVS were tightly overlapping to assemble a single cluster, the domain of ICVS in Ift88^f/f^;Isl1-Cre mutants was broader and, in some cases, formed two foci of cells (Figure 5D) Sox9-expressing cells accumulated in the proximal region of the outflow tract and assembled as a separate cluster from Isl1-expressing cells (n=4, Figure 5D), supporting the idea that there were two foci in the mutant ICVS. To understand the relationship between these two populations of cells within the ICVS, quantification of these cells (Isl1+ and Sox9+ cells) was performed. This showed that there were fewer cells expressing Isl1 (red) alone and more cells expressing solely Sox9 (green), or co-expressing both Sox9 and Isl1, in Ift88^f/f^;Isl1-Cre mutants compared to controls (n=6; Figure 5E). These data suggest premature differentiation of SHF progenitors to VIC in the ICVS of the Ift88^f/f^;Isl1-Cre mutants that led to fewer Isl1+ cells and more Sox9+ cells.

**Figure 5:**
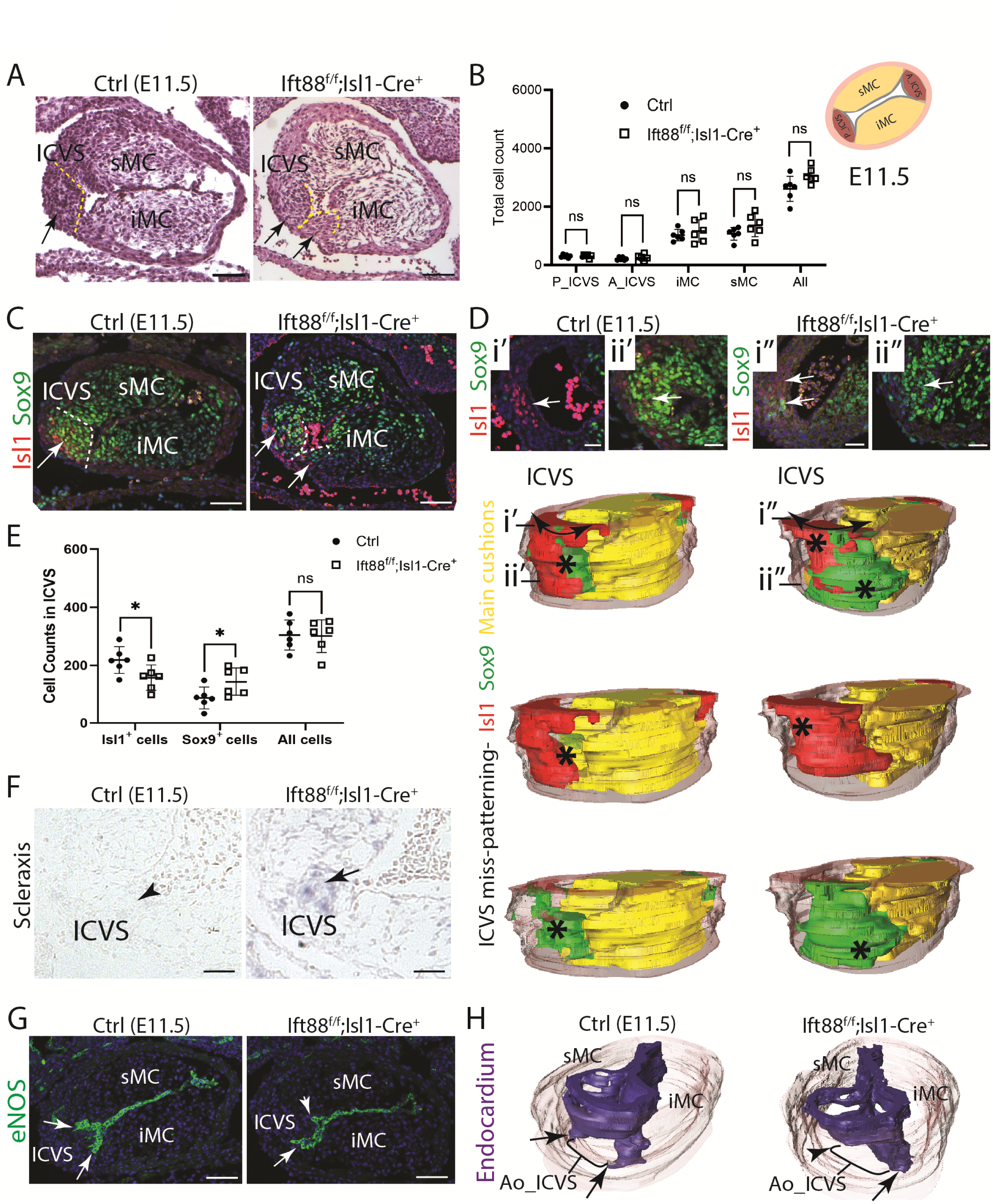
Premature differentiation of SHF cells correlates with abnormal ICVS formation in Ift88^f/f^;Isl1-Cre mutants during early development. **A)** Histological analysis of the outflow tract in E11.5 Ift88^f/f^;Isl1-Cre mutants (n=14), compared with their control littermates, revealed that although the ICVS was present in all cases, the mutant ICVS appeared to form two foci compared to the single focus in control littermates (12/14; arrows). **B)** Quantification of total cell counts in the ICVS and main cushions (superior sMC and inferior iMC) at E11.5 showed comparable overall counts between Ift88^f/f^;Isl1-Cre mutants and their littermate controls, n=6 for each genotype. Two-way ANOVA with Šídák test, Mean ± SD. ns: not significant. **C)** Matched immunofluorescence images for Isl1 (red; undifferentiated SHF cells) and Sox9 (green; partially differentiated cushion cells) confirmed the presence of two foci in the posterior ICVS (arrows in ii) of Ift88^f/f^;Isl1-Cre^+^ mutants. **D)** 3D reconstructions of serial immunofluorescent images of the outflow tract, labelling main cushions (yellow) and ICVS with Isl1 (red) or Sox9 (green) revealed that the ICVS in Ift88^f/f^;Isl1-Cre^+^ mutants was broader (double headed arrows), with Sox9-expressing cells appearing to assemble a separate focus from the Isl1+ cells (asterisks). The expression of both Isl1 (red) and Sox9 (green) at the distal region (arrows in i’’) and the sole proximal expansion of Sox9+ cells (arrow in ii’’) in Ift88^f/f^;Isl1-Cre^+^ mutants; support the idea that there are two foci forming in the mutant ICVS. **E)** Quantification of Sox9- and Isl1-expressing cells in the ICVS at E11.5 indicated significantly fewer Isl1^+^ cells, and more Sox9^+^ cells in the Ift88^f/f^;Isl1-Cre^+^ compared to that in control littermates, n=6 for each genotype. Two-way ANOVA with Šídák test, Mean ± SD. ns: not significant, *p<.05. **F)** In situ hybridisation showed that *Scleraxis* was highly expressed in the ICVS of 3/4 the Ift88^f/f^;Isl1-Cre^+^ mutants (arrow) at E11.5, whereas there was little or no expression of *Scleraxis* in any of the littermate controls at E11.5 (arrowhead, n=4 for each genotype). **G,H)** Labelling of the cushion/ICVS endocardium with eNOS (green) at E11.5 showed that the endocardium forms a regular arc over the aortic ICVS in the control heart (arrows), whereas it is foreshortened, reflecting the juxtaposition of the supernumerary ICVS with the adjacent cushion, in the Ift88^f/f^;Isl1-Cre^+^ mutant heart (arrowhead in G). This can also be seen in 3D reconstructions of the endocardium (H). ICVS, intercalated valve swellings; sMC, superior main cushion; iMC, inferior main cushion. Scale bar =100μm

In situ hybridisation for *Scleraxis*, which upregulates with VIC differentiation, was observed in the ICVS of 3 of 4 Ift88^f/f^;Isl1-Cre mutants at E11.5, whereas this was not apparent in any of the control littermates (Figure 5F), supporting the idea that there is premature differentiation of DD-SHF cells to VIC in Ift88^f/f^;Isl1-Cre embryos. There was no evidence of premature differentiation to other derivatives of the SHF, such as myocardium or smooth muscle cells, in the outflow region of Ift88^f/f^;Isl1-Cre mutants (Supplementary Figure 5 A,B). Altogether, this showed that the basis of the Ift88^f/f^;Isl1-Cre mutant phenotype at E11.5 was the formation of two foci within the ICVS, which most likely resulted from premature differentiation of SHF cells to VIC.

Immuno-labelling of the endocardium with eNOS highlighted the abnormal shape and position of the forming ICVS in the Ift88^f/f^;Isl1-Cre embryos at E11.5, revealing that whereas the endocardium covering the ICVS formed a regular arc over the ICVS in wild type embryos, this was more irregular in Ift88^f/f^;Isl1-Cre embryos, foreshortening towards the lumen in regions where there was a bulge of ICVS lateral to its normal position and was thus the adjacent ICVS and cushion were in close apposition (Figure 5 G-H). Examination of control embryos at this timepoint shows that the endocardium does not extend as far as the outflow wall and thus does not completely separate the ICVS and the main cushions (arrows in Figure 5G-H and supplementary Figure 5C); this region of continuity is expanded in the Ift88^f/f^;Isl1-Cre embryos with no caspase-3-labelled cells observed in this region (Supplementary Figure 5D). Thus, the foreshortened endocardium in the Ift88^f/f^;Isl1-Cre embryos reflects the abnormal bulging of the ICVS in this position rather than any specific abnormality of the endocardium.

At E12.5, in the six examined mutants, 2/6 had aortic valves with three leaflets. 3D reconstructions of these cases identified that although there were three leaflets, the posterior leaflet appeared slightly smaller than that in the controls (Supplementary Figure 6A). However, comparing the aortic valve leaflet volumes, based on data from the 3D reconstructions, between Ift88^f/f^;Isl1-Cre embryos and control littermates showed no significant differences between mutants and controls, suggested no changes in the size of leaflet primordia at this stage (Supplementary Figure 6B). In 1/6 mutants, the leaflet primordia were developmentally delayed, as their appearances were similar to those of E11.5 hearts, with bulky leaflet primordia and unseptated outflow tract (Supplementary Figure 6C). In the remaining 3/6 cases examined at E12.5, four leaflets were seen, with large well-formed left and right main leaflet primordia but two small leaflets found in the posterior leaflet position (arrows and arrowheads Figure 6 A,B). Furthermore, whilst the forming commissures in control aortic valves were positioned deep, close to the outflow tract wall, in the mutant quadricuspid valves they were shallow (Figure 6A and Supplementary Figure 6D), reflecting the foreshortened endocardium that was observed where the supernumerary bulges of ICVS abutted the main cushions at E11.5 (compare with Figure 5H). Thus, there was no endocardium between forming leaflets in these regions at either E11.5 or E12.5. No caspase-3-labelled cells were observed in the forming leaflets of wild type or mutant hearts at this or earlier timepoints (Supplementary Figure 5D, 6E), supporting the idea that there was no need for removal of endocardial cells from a remodelling fusion seam, as endocardial fusion between forming leaflets did not occur. 3D reconstructions of the aortic valve leaflets showed that the leaflets were grossly misshapen and confirmed the abnormal positioning of the commissures. (Figure 6B). To investigate if hyperplasia was causing the supernumerary leaflets, quantification of cell numbers in the valve leaflet primordia (excluding the developmentally delayed mutant) revealed that there were no significant differences in the total number of cells within the forming valve leaflets between Ift88^f/f^;Isl1-Cre embryos and control littermates at E12.5 (Figure 6C). To clarify, if there were two posterior leaflet primordia in Ift88^f/f^;Isl1-Cre mutants they were added together and presented as a separate group for these analyses so that they could be compared directly with controls. This suggested that whilst there were a subset of Ift88^f/f^;Isl1-Cre mutants with supernumerary leaflets in the position of the posterior leaflet, this was not due to increase in cell number within the leaflet primordia. By E13.5, the arterial valve primordia were beginning to remodel in control hearts, with sinuses appearing between the leaflets and the vessel wall. As before, some Ift88^f/f^;Isl1-Cre mutants were presented with three aortic leaflets (n=3/7). However, while a representative 3D reconstruction of these mutant leaflets showed that the posterior leaflet appeared slightly smaller compared to that in controls (Supplementary Figure 7A), quantification of leaflet volume and the total cell number for these mutants revealed no significant differences compared to controls (Figure 6F and Supplementary Figure 7B). In 2/7 mutants, and similar to the observed case at E12.5, the valve leaflets were developmentally delayed and appeared similar to those of E11.5 hearts with bulky cushion primordia and upseptated outflow tract (Supplementary Figure 7C). The remaining examined (2/7) mutants had large left and right leaflets, but only one small posterior leaflet (Figure 6D). 3D reconstructions of these valve leaflets (Figure 6E) confirmed the presence of a small posterior leaflet but showed an enlarged right leaflet that suggesting remodelling and fusion of the right main cushion primordia with one of the additional posterior primordia observed at E12.5. Interestingly, quantification of cells within the forming leaflets and their volume (for these two cases) suggested that although the total cell number and total leaflet volume for all leaflets together did not change, the posterior leaflet had fewer cells and reduced leaflet volume while the right leaflet had more cells and increased leaflet volume in the Ift88^f/f^;Isl1-Cre mutants compared to controls (Figure 6F and Supplementary Figure 7B). This suggested that the changes in the architecture of the leaflets were not due to hypoplasia, but rather to altered cell distribution within the leaflets.

**Figure 6:**
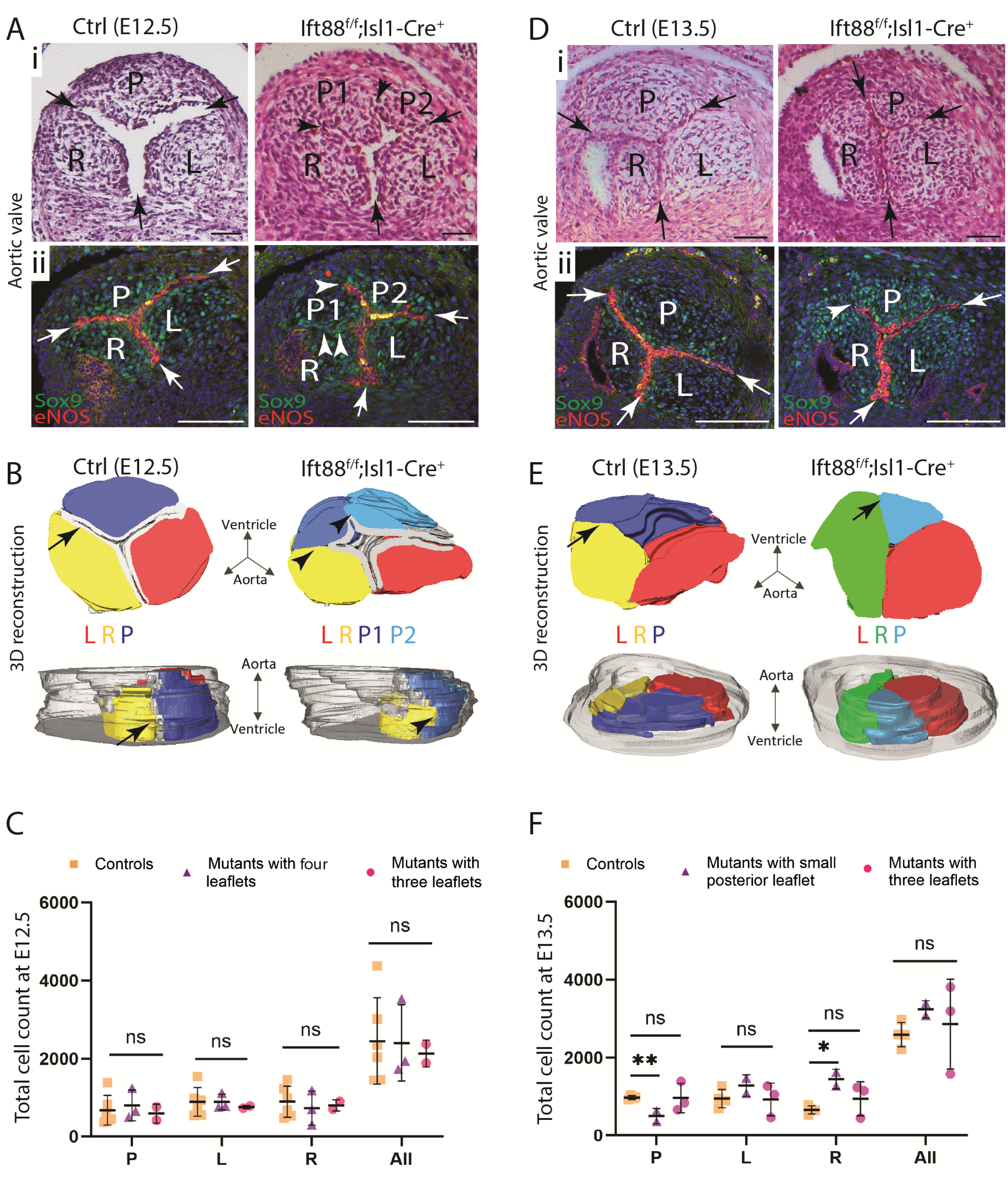
BAV in the Ift88^f/f^;Isl1-Cre mutants caused by remodelling of supernumerary leaflet primordia. **A)** Histological analysis of the aortic valve leaflet primordia at E12.5 revealed the presence of four primordia in 3/6 Ift88^f/f^;Isl1-Cre mutants, compared to the usual three primordia seen in control littermates. (i) Arrows point to the commissures where adjacent leaflets attach to each other at the aortic wall, whereas arrowheads indicate displacement of commissures from the aortic wall towards the lumen. (ii) Immunolabelling of cushion mesenchyme (labelled by Sox9; green) and the endocardium overlying each leaflet primordium (labelled by eNOS; red) shows that all three commissures were close to the arterial wall in controls (arrows in ii) whereas at least one commissure was displaced towards the lumen in Ift88^f/f^;Isl1-Cre^+^ mutants (arrowheads in ii; n=3/3. **B)** 3D reconstructions, produced from anti-Sox9 and anti-eNOS immunofluorescence images, revealed the extent of the endocardium in each case (pale grey), relative to each of the primordia in the controls (P is blue, R is yellow, L is red) and the Ift88^f/f^;Isl1-Cre^+^ mutants (two P are different shades of blue, R is yellow, L is red). In the Ift88^ff^;Isl1-Cre mutants, the commissures are close to the lumen between the R and P1 primordium (arrowheads), with continuity in the mesenchyme between these conjoined primordia (arrowheads), n=3/6. **C)** Total cell counts from each of the aortic leaflet primordium revealed no significant differences between Ift88^f/f^;Isl1-Cre^+^ mutants and their littermate controls at E12.5. Where there was the appearance of two P primordia in the mutants, these were counted together (purple triangles) to allow comparison with controls. Mutants with three leaflet primordia were presented as pink circles. Two-way ANOVA with Šídák test, Mean ± SD. ns: not significant. **D)** Histological analysis of the aortic valve leaflets at E13.5 showed that the primordia were remodelling and could now be described as leaflets. A total of three leaflets, but with a small P leaflet could be seen in 2 of 7 mutants (arrows in i pointing at the commissures). Immunolabelling of mesenchymal cells with Sox9 (green) and the endocardium overlying each leaflet primordium (labelled by eNOS, red) again showed displacement of the commissure between P and the adjacent R leaflet (arrowhead in ii) towards the lumen. **E)** 3D reconstructions, produced from anti-Sox9 immunofluorescence images without labelling of the endocardium, showed a small P leaflet (light blue) with a remodelling R leaflet (green) appearing to emerge more centrally (arrows) in Ift88^f/f^;Isl1-Cre mutants compared to that in controls (P is blue, R is yellow and L is red), n=2/7. **F)** Total cell counts from each aortic leaflet revealed significant differences in the posterior and right leaflets between Ift88^f/f^;Isl1-Cre mutants and their littermate controls, but not in the overall cell counts for all three leaflets at E13.5. Mutants with small P leaflets were presented as purple triangles, whereas mutants with three leaflets were presented as pink circles. Two-way ANOVA with Šídák test, Mean ± SD. ns: not significant, **p=.006, *p=.024. L, left leaflet; R, right leaflet; P, posterior leaflet; P1 and P2, super-numerary leaflets in the position of the usual posterior leaflet. Scale bar =100µm

Together, these data suggest that a broadened and prematurely differentiating ICVS at E11.5 in Ift88^f/f^;Isl1-Cre mutants leads to the formation of two leaflet primordia in the position of the usual posterior leaflet at E12.5, pushed against the leaflet primordia from the adjacent main cushions. These ICVS-derived primordia, with some variation, merged with the adjacent right (and possibly left) leaflets between E13.5 and E15.5, resulting in BAV.

### Hedgehog and Wnt signalling are not altered in ICVS of Ift88^f/f^;Isl1-Cre mutants

To begin to uncover the mechanism of premature differentiation of the ICVS, we first investigated potential disruption to Hedgehog signalling as this is a recognised consequence of cilia loss ^36^. We therefore looked at the expression of Ptch1 and Gli3 in the developing ICVS at E11.5, when the aortic valve leaflet defects in the Ift88^f/f^;Isl1-Cre mutants first become apparent. RNAscope showed that both Ptch1 and Gli3 were expressed at similar levels in the ICVS at this time point (Supplementary Figure 8 A,B; n=3 for each genotype). Earlier analysis of the outflow region and immediately adjacent SHF in Ift88^f/f^;Isl1-Cre mutants and control littermates showed that although there was upregulation of Ptch1 in the pharyngeal and paraxial mesoderm (n=4 for each genotype), there were no changes in the distal OFT or the adjacent region of the SHF for Ptch1 or Gli3 (Supplementary Figure 8C).

Canonical Wnt signalling can also occur at the cilium^37^ so we examined the expression of Lef1, a readout of canonical Wnt signalling. Interestingly, whereas Lef1 was not expressed in the outflow tract at E10.5 (Supplementary Figure 8D), it specifically localised to the ICVS at E11.5 in both control and Ift88^f/f^;Isl1-Cre embryos (Supplementary Figure 8E). However, there were no differences in the numbers of Lef1-expressing cells between controls and mutants (n=4, Supplementary Figure 8F). We also looked at the expression of Axin2, as an additional readout of canonical Wnt signalling, but it did not localise to the developing ICVS (Supplementary Figure 8G). We therefore conclude there is no evidence that Hh or canonical Wnt signalling is impacted in the ICVS of Ift88^f/f^;Isl1-Cre embryos.

### Notch signalling is disrupted in the ICVS of Ift88^f/f^;Isl1-Cre mutants

Notch signalling plays a major role in the development of the outflow tract ^38,39^. More specifically, loss of Notch signalling in the DD-SHF lineage results in loss of the posterior leaflet and BAV ^18^. Thus, we next examined the impact of loss of cilia on Notch signalling in the developing ICVS in Ift88^f/f^;Isl1-Cre mutants.

Using immunofluorescence, we examined the expression of active Notch1 (N1ICD), Notch2, Notch3 and Jag1 in the developing ICVS at E10.5 and E11.5. Whereas Notch1ICD and Jag1 were specifically localised to the developing ICVS both at E10.5 and E11.5 (Figure 7A) neither Notch2 nor Notch3 did at E10.5 or E11.5 (Supplementary Figure 9A). Previous studies have shown that both Notch1ICD and Jag1 localise to Isl1-expressing SHF cells in the developing ICVS [18] (Figure 7A-B). As shown earlier (Figure 7C), these Isl1-expressing cells are ciliated.

**Figure 7:**
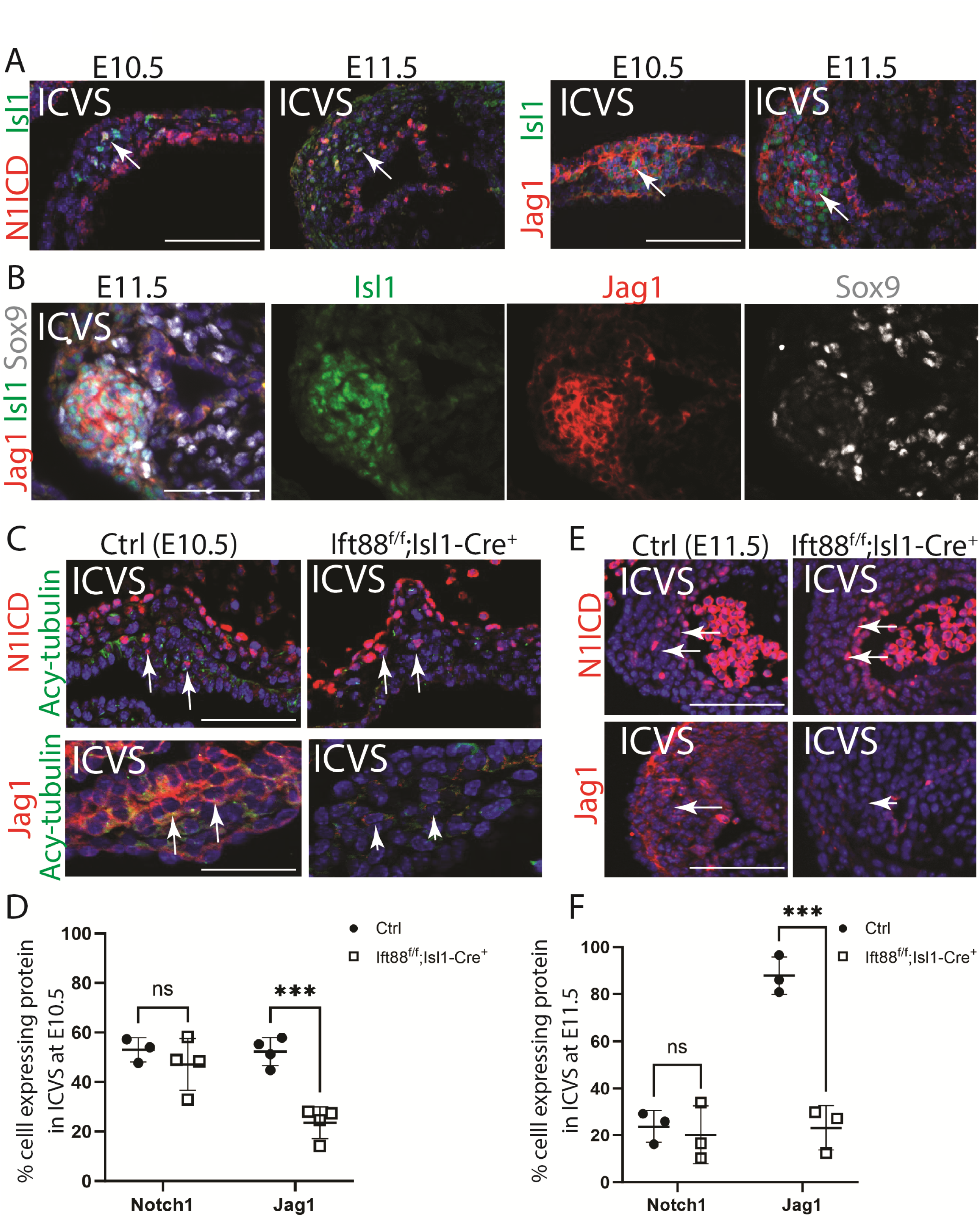
Disruption of Notch signalling pathway in the outflow tract of Ift88^f/f^;Isl1-Cre^+^ mutants. **A)** Active Notch1 (N1ICD, red) was abundant in the ICVS at E10.5, co-localising to Isl1-expressing cells (green), but was found in only a few ICVS cells by E11.5 (arrows). Jag1 (red) localised to Isl1-expressing cells (green) in the ICVS at both E10.5 and E11.5 (arrows). **B)** The expression of Jag1 (red) is largely restricted to undifferentiated Isl1-expressing cells (green) in the ICVS at E11.5. **C-E)** At E10.5 and E11.5, N1ICD in the ICVS was comparable between Ift88^f/f^;Isl1-Cre mutants and their littermate controls (arrows). In contrast, Jag1 showed reduced expression in the ICVS (arrowheads) compared to littermate controls (arrows) at both stages n=3/3 (C,E). Quantification confirmed that whilst for N1ICD there were no significant differences between mutants and control littermates in the number of positive cells expressing the receptor, the number of positive cells expressing Jag1 was dramatically reduced in the mutants compared to control littermates at E10.5 and E11.5 (D,F). Two-way ANOVA with Šídák test, Mean ± SD. ns: not significant, ***p<.001. ICVS, intercalated valve swellings. Scale bar =100µm.

Turning to the mutant line, we observed that whereas N1ICD expression appeared unaltered in the ICVS of Ift88^f/f^;Isl1-Cre embryos at E10.5, the number of cells expressing Jag1 was significantly reduced in the mutant ICVS at both E10.5 and E11.5 (Figure 7 C-F), correlating with the reduced Isl1 expression and loss of cilia (see Figure 4G). This reduced Jag1 expression extended to the anterior and posterior region of the dorsal pericardial wall and distal walls of the outflow tract at E10.5 (Supplementary Figure 9B). Importantly, this abnormal expression occurred before the outflow tract appeared abnormal in the Ift88^f/f^;Isl1-Cre embryos and thus could be contributing to the later abnormalities. Hes1 and Hey2 are mediators of Notch signalling expressed in the developing outflow tract ^40^. Using RNAscope, we analysed the expression of Hes1 and Hey2 in the Ift88^f/f^;Isl1-Cre mutants compared to their littermate controls at E10.5. This revealed a marked reduction in the expression of Hes1 in the dorsal pericardial wall and distal outflow tract, and Hey2 in the endothelial cells of the outflow tract in the mutants (n=3/4; Supplementary Figure 9 C-D). Similar levels of Hes1 in the neural tube and Hey2 in ventricle were found in Ift88^f/f^;Isl1-Cre mutants and controls (Supplementary Figure 9E), suggesting the reductions were specific to the Isl1-Cre expressing regions of the embryo. Taken together, this suggests that a loss of Jag1-dependent signalling may lead to the premature differentiation of cells in the posterior ICVS that, following primordia remodelling results in smaller or absent posterior leaflet.

To test the idea that Jag1 signalling in the SHF is required for normal development of the ICVS, we knocked it out in the Isl1 lineage. Examination of litters at E14.5-E16.5 revealed that 7 of 10 (70%) had abnormalities of the arterial valves, which included BAV (Figure 8A, Supplementary Figure 10A). We then examined the developing ICVS at E11.5 in the Jag1^f/f^;Isl1-Cre mutants, to see if there was evidence for a broadened ICVS and/or premature differentiation of the SHF. Similar to the Ift88^f/f^;Isl1-Cre mutants, the ICVS of Jag1^f/f^;Isl1-Cre mutants had fewer Isl1+ cells and more dual-labelled Isl1+/Sox9+ and Sox9+ cells within the ICVS (Figure 8B, n=3). Therefore, the observation of premature differentiation (fewer Isl1+ cells, more Sox9+ cells) in the Jag1^f/f^;Isl1-Cre mutants compared to controls suggests the involvement of Jag1 in regulating the differentiation of Isl1-positive cells within the ICVS to VIC during valve development. The implication is that Jag1, likely acting through cilia, is crucial for the proper timing and progression of this differentiation process.

**Figure 8:**
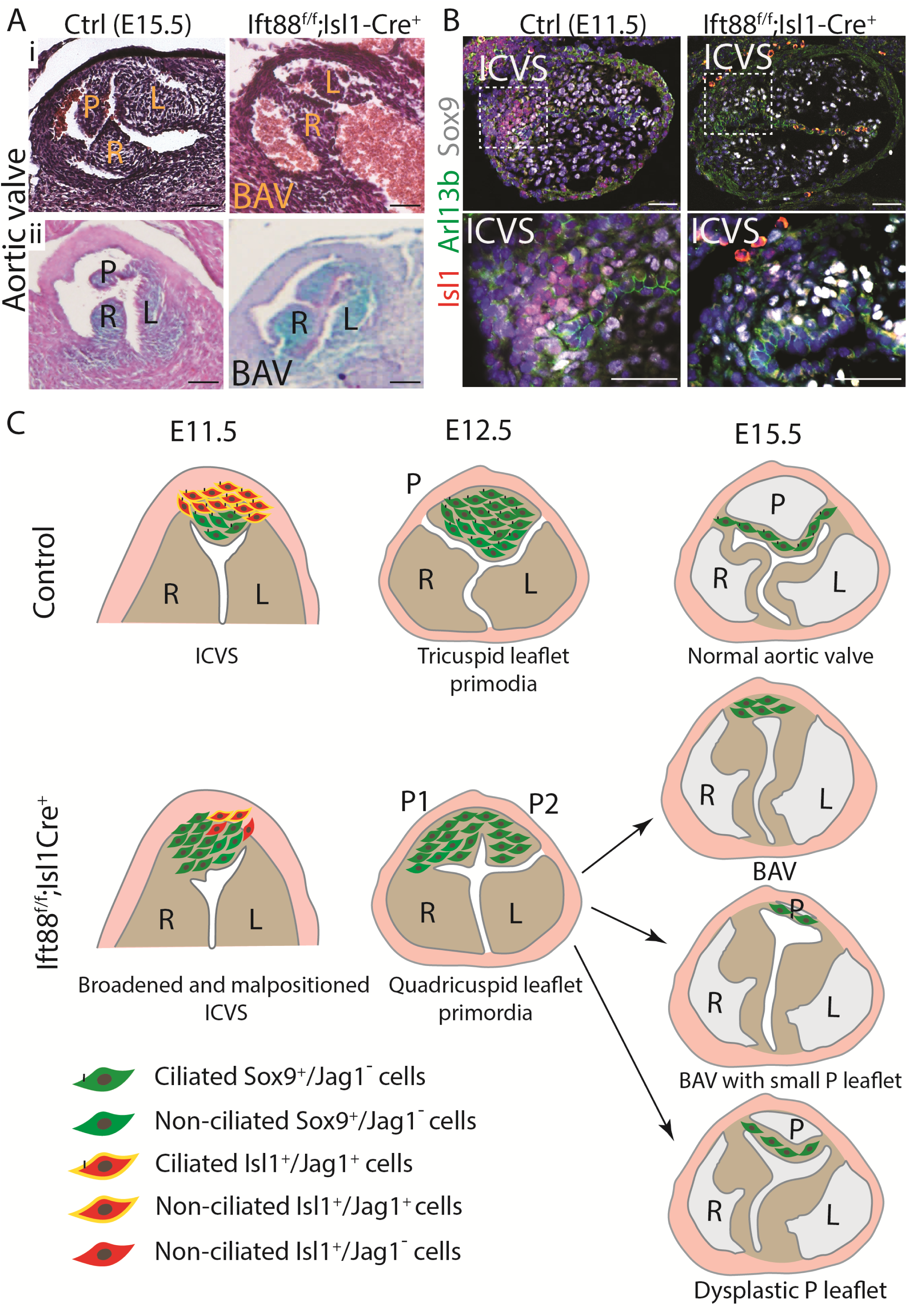
Aortic valve anomalies associated with premature differentiation of SHF cells in the ICVS of Jag1^f/f^;Isl1-Cre mutants. **A)** Arterial valve defects, including BAV, in E15.5 Jag1^f/f^;Isl1-Cre mutants compared to littermate controls detected with H&E staining (i) and Alcian Blue (ii). **B)** Immunofluorescence for Isl1 (red), Sox9 (white) and Arl13b (green) reveals that there are fewer Isl1+ cells, but more Sox9+ cells in the ICVS of the Jag1^f/f^;Isl1-Cre mutant compared to control littermates, n=3. **C)** Cartoon model illustrating how loss of primary cilia from Isl1-expressing cells disrupts aortic valve development and results in BAV. In controls, ICVS cells differentiate from Isl1-expressing progenitors (red) to VIC expressing Sox9 (green) as they loss Jag1 expression (yellow) between E11.5-E12.5. They carry primary cilia during this period. Conversely, in mutants, when primary cilia are lost from the undifferentiated SHF cells (red) they lose their expression of Jag1 (yellow) and differentiate prematurely to Sox9 (green). This results in a broadened ICVS that merges with the main cushions leading to the appearance of BAV as the primordia remodel. L, left leaflet; R, right leaflet; P, posterior leaflet; ICVS, intercalated valve swellings. Scale bar =100µm

## Discussion

In this study, we have shown that primary cilia were found in the majority of cells in the early developing outflow tract but by E12.5, less than 50% were ciliated. Cells dissemble their primary cilium as they divide and reassemble them once cell division is complete ^36^. However, this ciliation event did not correlate with increases in proliferation of the forming valve leaflets. Instead, the decline in the percentage of ciliated mesenchymal cells in the forming valve leaflets correlated with the expression of *Scleraxis*, which is required for cushion cell differentiation ^32^. Cilia loss also correlated with differentiation of cardiomyocytes and smooth muscle cells in the developing outflow tract, suggesting that this process may be related to differentiation of at least some cell types. In support of a role for cilia in regulating cell differentiation, we observed premature differentiation of SHF-derived cells in the forming ICVS in Ift88^f/f^;Isl1-Cre mutants, ultimately leading to BAV, supporting the idea that cilia may be co-ordinating differentiation in the normal situation. It is known that some VIC are ciliated in the adult heart, and that their loss, via deletion of Ift88, results in enlarged, myxomatous valves ^11^. Thus, it seems likely that cilia play roles in mature valves, potentially in maintaining valve homeostasis.

In our hands, deletion of Ift88 from NCC or endocardial cells did not result in any outflow defects, despite effects on other parts of the embryo including other structures within the heart. With respect to NCC, our data is in agreement with previous authors, who reported severe craniofacial defects when Ift88 was deleted using Wnt1-Cre, but without mention of the outflow tract ^33^. However, our findings are at odds with the findings of ^13^ who found a high incidence of BAV when Ift88 was knocked out in the endocardium using Nfatc1-Cre. As we observed reduced numbers of ciliated cells in the Tie2-Cre-positive cushion cells in the Ift88f;Tie2-Cre mutant fetuses, and other heart malformations were observed, it seems unlikely that the knockout was inefficient. It may be, however, that as we used different Cre drivers, differences in the penetrance or timing of Ift88 deletion explains the discrepancy. We did see arterial valve defects, however, when Ift88 was deleted using Tnnt2-Cre, which as well as targeting the myocardium, labels directly-differentiating SHF (DD-SHF) that make a major contribution to the ICVS, with a more minor contribution to the right and left leaflets ^18^. Loss of cilia in this population gave rise to defects in the ventricular myocardium but also dysplastic arterial valve leaflets. Perhaps surprisingly, all three leaflets appeared to be affected equally, perhaps suggesting a role for cilia in the Tnnt2-Cre+ cells that are found in all three leaflets ^18^. Alternatively, there may be effects on the arterial valves secondary to gross abnormalities in the ventricular myocardium in the Ift88f;Tnnt2-Cre mutants.

When Ift88 was deleted using the SHF-specific driver, Isl1-Cre, arterial valve defects were seen in more than half of the mutants. The observation that neither ourselves, nor Burns et al., 2019 ^12^, observed arterial valve defects following deletion of Ift88 with Mef2c-AHF-Cre, could be explained by a number of possibilities. Firstly, the Isl1-Cre allele [25] that we use is a knock-in, meaning that mice carrying this allele are haplo-insufficient for Isl1. As it is Isl1-expressing cells in the ICVS that we believe are critical to the development of the phenotype in the I Ift88^f/f^;Isl1-Cre mutants, reduced levels of Isl1 could have an impact. Arguing against this, we did not observe reduced expression of Isl1 in the Isl1-Cre+ hearts and the Isl1-Cre animals are not reported to have a phenotype [25]. Alternatively, the observation of BAV in the Isl1-Cre, but not the Mef2c-AHF-Cre mutants, could reflect a difference in the precise cells labelled by each Cre driver, related to timing or precise pattern of expression. Close examination of the expression domains showed that at E11.5 and E12.5, there was a population of cells in the valve endocardium and underlying mesenchyme of the aortic ICVS that was labelled by Isl1-Cre, but not by Mef2c-AHF-Cre. Whether either of these differences is enough to explain the difference between the phenotypes when Ift88 is knocked out using the two Cre drivers, or whether it relates to earlier differences in pattern or timing of activation, remains to be shown. The observation that Isl1-Cre labels a sub-population of NCC in the outflow cushions ^35^ seems unlikely to be the explanation, as loss of all cilia from all cells using Wnt1-Cre as a driver did not result in outflow abnormalities.

There are several suggested mechanisms for how BAV may arise ^41,42^. Moreover, it has been suggested that there may be an ontological link between quadricuspid and bicuspid leaflets. Clinically, quadricuspid aortic valve (QAV) is found in up to 0.04% of the population ^43^, so is much less common than BAV. Interestingly, QAV and BAV are both reported in a strain of Syrian hamsters ^44^. Moreover, we have previously described the presence of two leaflet primordia in the position of the normal posterior leaflet in the ROCKDN;Wnt1-Cre mouse mutant, where both BAV and QAV were observed at E15.5 ^45^. Here in Ift88^f/f^;Isl1-Cre mutants, although we have not observed any cases with QAV at E15.5, a subset of Ift88^f/f^;Isl1-Cre had the appearance of quadricuspid leaflet primordia at early stages. At E11.5, the ICVS of these Ift88^f/f^;Isl1-Cre were broader with premature differentiation of Isl1^+^ progenitors to Sox9-expressing VIC that formed two separated foci within the region where the posterior leaflet forms. Presumably these two foci in the ICVS were the precursors of the two small posterior leaflet primordia that were observed by E12.5. Close apposition of the ICVS and the main cushions, prior to the formation of endocardium between them, then results in the appearance of four forming leaflets at the lumenal surface of the forming valve, but continuity of mesenchyme between them more laterally. As development proceeds, at E12.5 and E13.5, there is thus no need to remodel an endocardial fusion seam, as such a fusion seam never existed, with the fusion occurring prior to the normal extension of the endocardium towards the walls of the arterial roots to cover the entirely of the forming valve leaflet. In the healthy developing aortic valve, the leaflet primordia are arranged in close proximity and occlude the outflow tract between heart beats ^46^. It seems likely that that the presence of an additional primordium leads to excessive apposition between adjacent primordia, resulting in merging of adjacent primordia as the leaflets continue to remodel leading to BAV. It is unclear whether a broadened ICVS is a common model for the formation of quadricuspid leaflets, but the most common patterns of QAV in human have supernumerary leaflets in the position of the normal posterior leaflet ^47–49^, similar to what we observe. Thus, a mechanism where a broadened ICVS leads to the formation of two leaflets in the position of the usual posterior leaflet, may be clinically relevant.

A number of different signalling pathways rely on cilia to transduce the signal from the cell surface to the nucleus, the best known of these is the Hedgehog (Hh) pathway^50^. Sonic hedgehog (Shh) signalling is known to be important for outflow tract development ^51^ and loss of cilia abrogates Shh signalling ^50^. However, there do not appear to be any direct associations in the published literature between Shh signalling and arterial valve development, although there is a link between Desert hedgehog (Dhh) signalling and development of the mitral valve ^10^. Surprisingly, despite the dramatic reduction in cilia numbers and length we observed in Ift88^f/f^;Isl1-Cre mutants compared to their control littermates, we did not find reproducible differences in the levels or distribution of the Shh signalling components Ptch1 and Gli3 in the ICVS or outflow tract, although Ptch1 was reproducibly upregulated in the pharyngeal and paraxial mesoderm in the mutants. Canonical Wnt signalling has also been associated with cilia [36] and has been linked to valve development ^52^. Despite this, and the specific expression of Lef1, a canonical Wnt mediator, in the developing ICVS, we did not find any differences in the numbers of cells expressing Lef1 in the Ift88^f/f^;Isl1-Cre embryos; other mediators we tested were not expressed in the developing ICVS. Thus, although not exhaustive, we could find no evidence that canonical Wnt signalling was altered by the loss of cilia in our study.

In contrast to the data for Hh and Wnt signalling, we saw a significant reduction in the Notch ligand Jag1 in the ICVS at E11.5 and in the distal outflow tract and the adjacent pericardial wall at E10.5. Hes1 and Hey2, downstream mediators of Notch signalling, were also downregulated in the outflow region at E10.5. Thus, we suggest a mechanism where loss of cilia in the cells of the forming ICVS leads to their premature differentiation and formation of two primordia in the position of the usual posterior leaflet. Although not as well described as Hh, there is significant evidence for a link between cilia and Notch signalling (reviewed in ^36,53,54^ and loss of cilia has been associated with abrogation of Notch signalling in a number of scenarios ^55,56^. In the outflow tract, loss of cilia in Ift88^f/f^;Isl1-Cre embryos resulted in a reduction in the levels of Jag1, Hes1 and Hey2, all of which have been shown to be important for outflow tract development, and specifically in the case of Jag1, the arterial valves ^18,57–59^. Indeed, conditional deletion of Jag1/Jag2 in DD-SHF cells in the ICVS, using Tnnt2-Cre, resulted in BAV characterised by absence of the posterior leaflet, similar to what we describe for the Ift88^f/f^;Isl1-Cre mutants. To add to this, we now show that deletion of Jag1 in the SHF, using Isl1-Cre as a driver, results in very similar arterial valve defects as when cilia are removed from the SHF-derived cells in the Ift88^f/f^;Isl1-Cre embryos. Moreover, the mechanism appears to be similar, as in both cases there was a broadened ICVS and premature differentiation of Isl1-expressing SHF cells to Sox9-expressing VIC at E11.5. This provides strong supporting evidence for the idea that Jag1 signalling is disrupted by loss of cilia in the SHF, and this impacts directly on the developing ICVS through premature differentiation, resulting in BAV (Figure 8C).

Large scale ENU mouse screens for mutants with CHDs have suggested a high burden of cilia related genes ^6^. However, whilst genomic studies have suggested that variants in cilial genes are enriched in rare familial cases, this is not the case for isolated de novo cases of CHD ^7,8^, raising the possibility that the interpretation of the mouse data was biased by the extensive delineation of the ciliome and the associated high number of Gene Ontology terms relating to cilia. Notably, almost 1000 genes have been associated with cilia ^60^, comparable to the 1200 genes reported to be expressed in mitochondria ^61^, another well studied organelle. Another possibility is that disruption of cilial genes may allow intrauterine development in mouse but may be lethal early in development in human, and hence never be encountered in human medical practice. It is important to recognise that the malformations seen in the Ift88^ff^;Tie2-Cre and Ift88^ff^;Tnnt2-Cre fetuses included major disruptions to myocardial architecture that may well be embryonic lethal during human pregnancy. In contrast, the malformations found in Ift88^ff^;Isl1-Cre, principally, BAV, are compatible with survival in utero and hence the case for a role for cilia in valve malformations, especially the aortic valve, is well made. Thus, early developmental loss due to non-survivable cardiac malformations may be a factor influencing the paucity of heart malformations in ciliopathy syndromes.

## Funding

We would like to thank the British Heart Foundation grants (RG/12/15/29935, RG/19/2/34256) and MRC National Mouse Genetics Network (MC_PC_21044) for funding this project. The Jag1^f/f^;Isl1-Cre study was supported by the Czech Science Foundation (24-12330K to HK).

## Author Contribution Statement

Conceptualization: A.A, B.C., D.J.H.; Methodology: A.A., L.E, B.C., D.J.H.; Software: A.A, D.J.H.; Validation: A.A, D.J.H.; Formal analysis: All authors; Investigation: A.A, L.E., C.J.D, B.C., D.J.H.; Data curation: All authors; Writing - original draft: A.A, D.J.H.; Writing - review & editing: All authors; Visualization: A.A., D.J.H.

## Conflict of interest

The authors have declared that no competing interests exist.

## Acknowledgements

We would like to thank the Comparative Biology Centre at Newcastle University for their support for animal studies.

## Notes

### Competing Interest Statement

The authors have declared no competing interest.

