## Supplementary figures for "Remodelling of supernumerary leaflet primordia leads to bicuspid aortic valve (BAV) caused by loss of primary cilia"

### Supplementary Figure 1

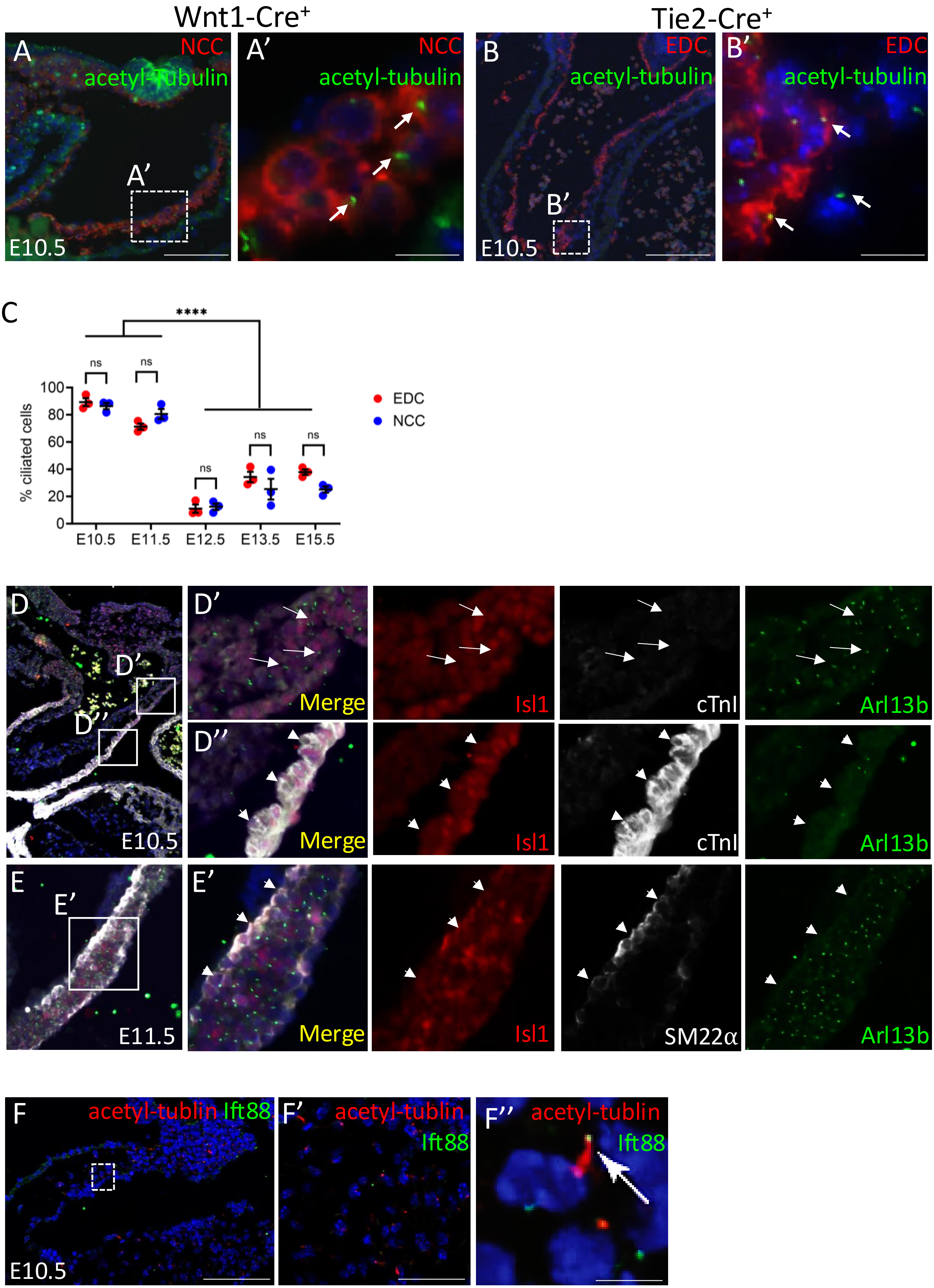

Supplementary Figure 2

A

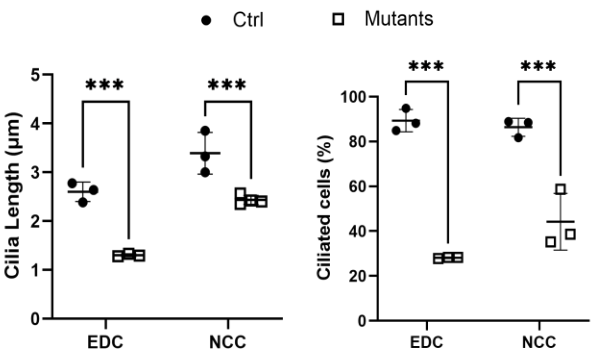


B

| | Mutant | Total | Survival % | Expected % |
|---|---|---|---|---|
| Ift88$^{f/+}$; Wnt1-cre | 3 | 33 | 8.6%* | 12.5% |
| Ift88$^{f/f}$ ; Wnt1-Cre | 5 | 27 | 18.5%** | 25% |
| Ift88$^{f/f}$; Tie2-Cre | 20 | 69 | 29% | 25% |
| Ift88$^{f/f}$; Tnnt2-Cre | 9 | 60 | 21% | 25% |

Chi squared equal 0.390* and 0.605**

C

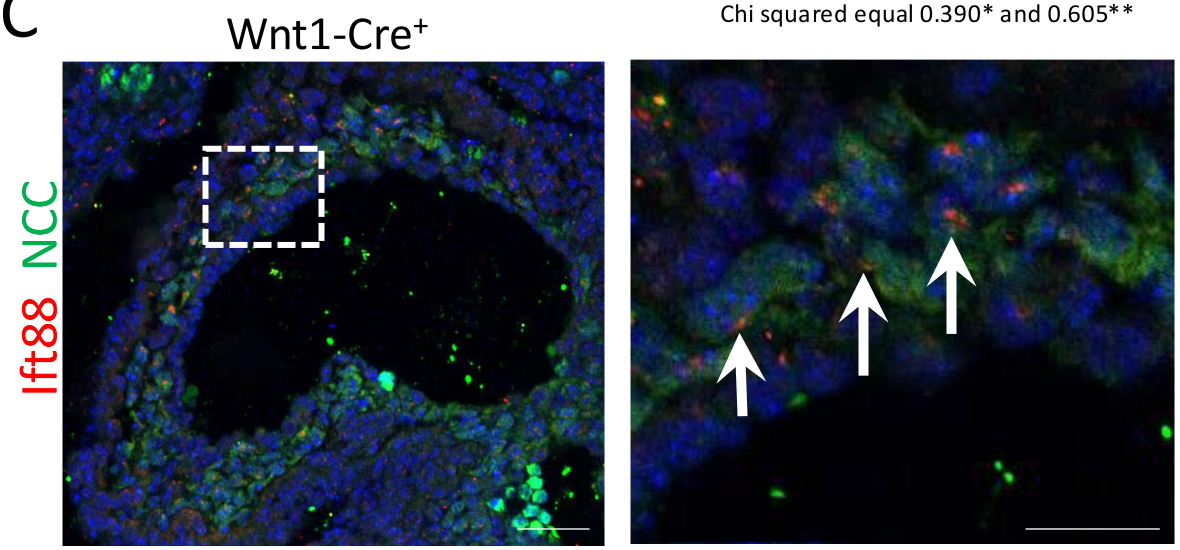


D

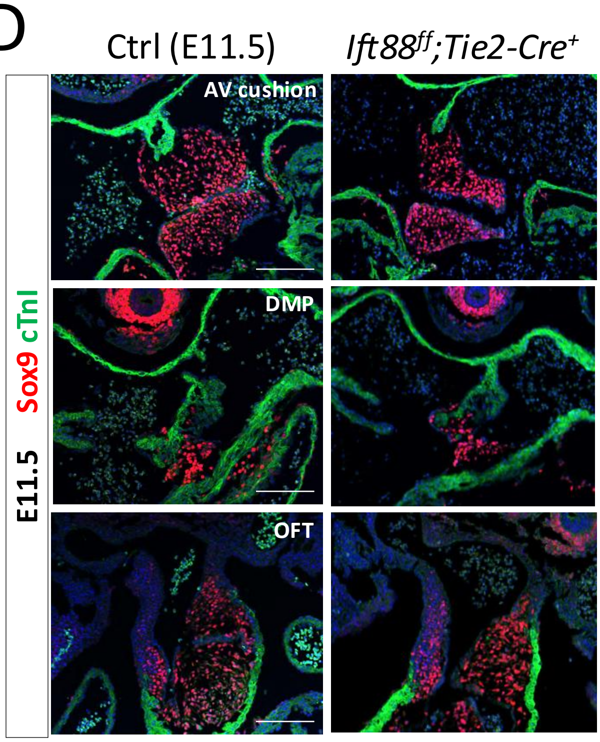


E

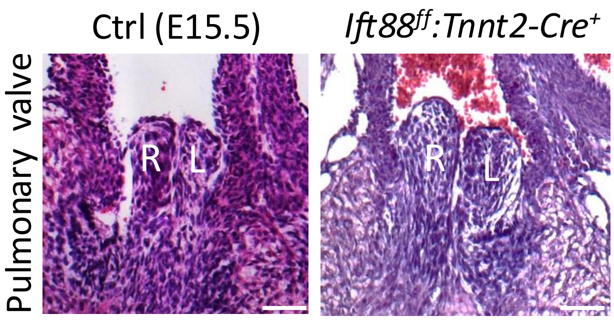

### Supplementary Figure 3

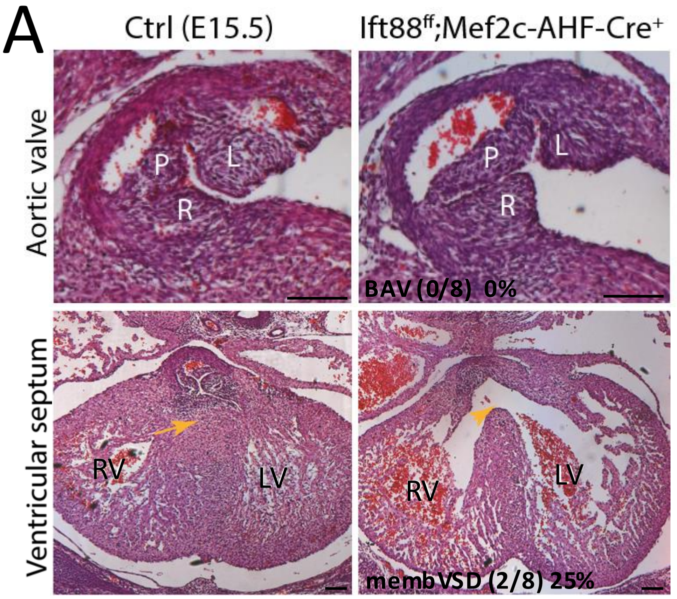


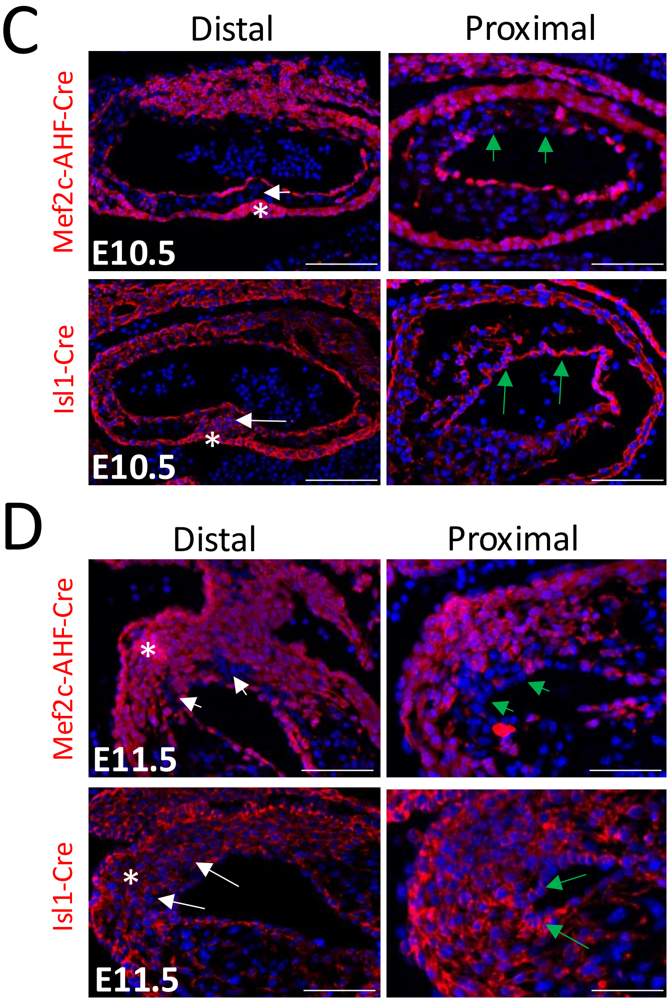


B

| | Mutant | Total | Survival % | Expected % |
|---|---|---|---|---|
| Ift88$^{f/f}$; Mef2c-AHF-Cre | 11 | 34 | 32%* | 25% |
| Ift88$^{f/f}$; Isl1-Cre | 19 | 71 | 27% | 25% |

*Chi squared equals 0.980

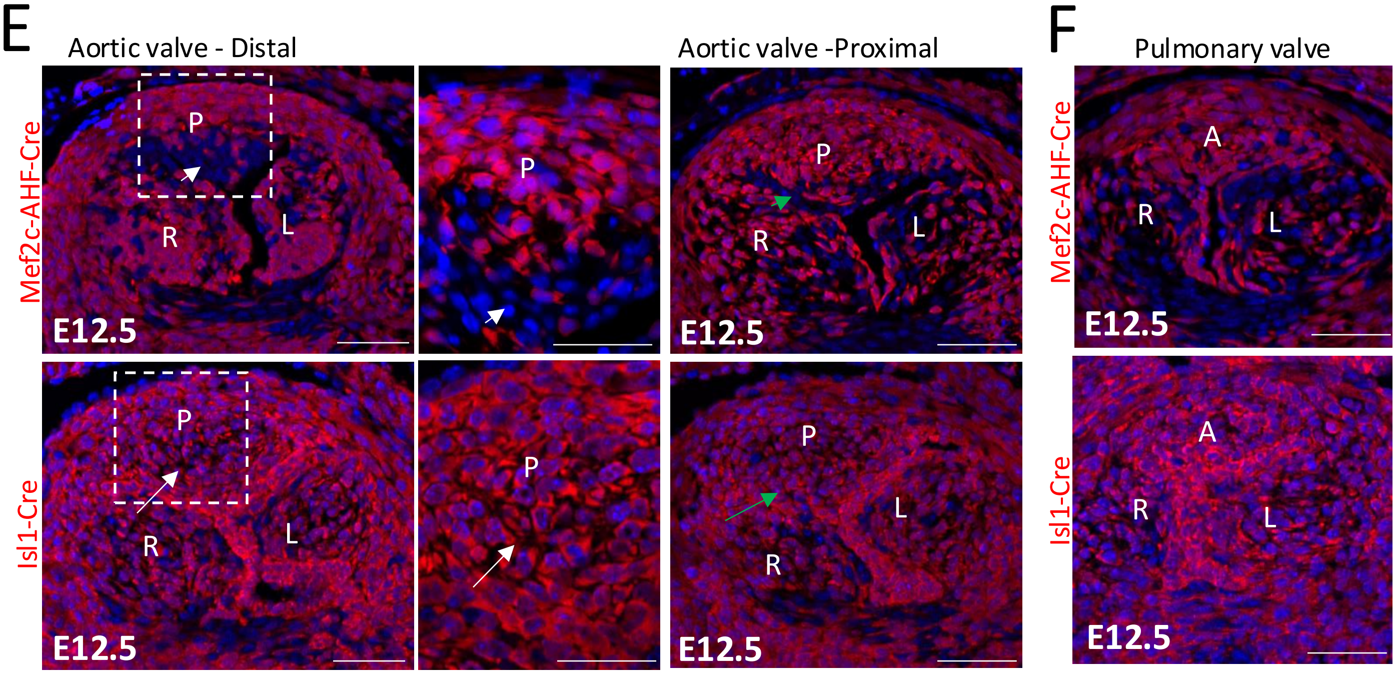

Supplementary Figure 4

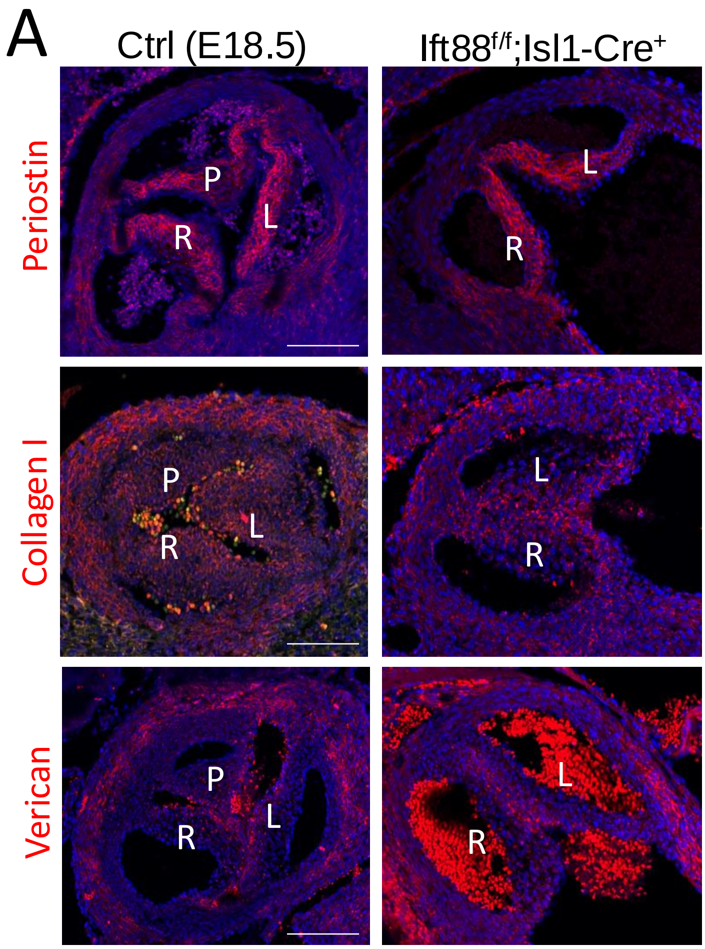

Supplementary Figure 5

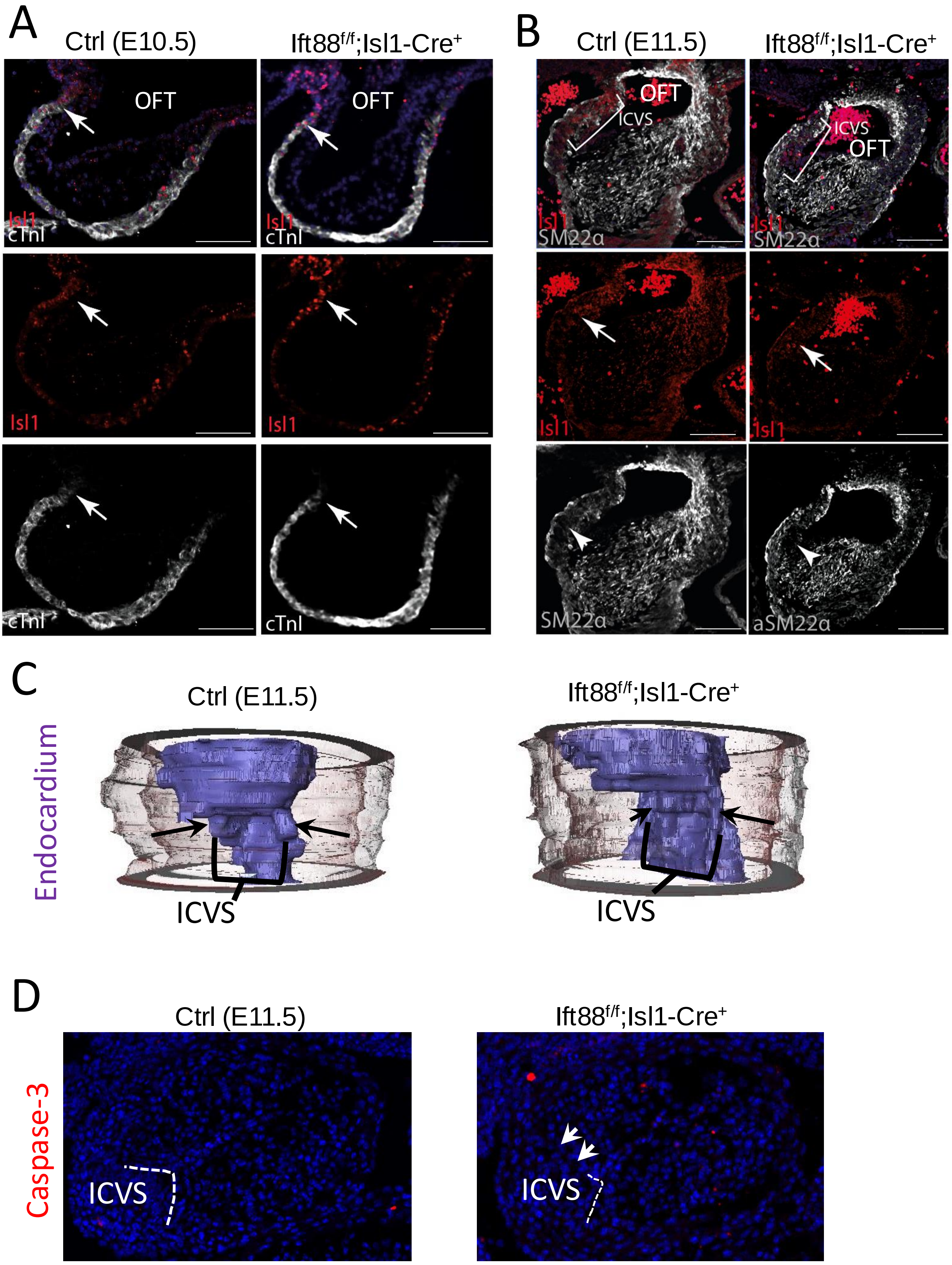

Supplementary Figure 6

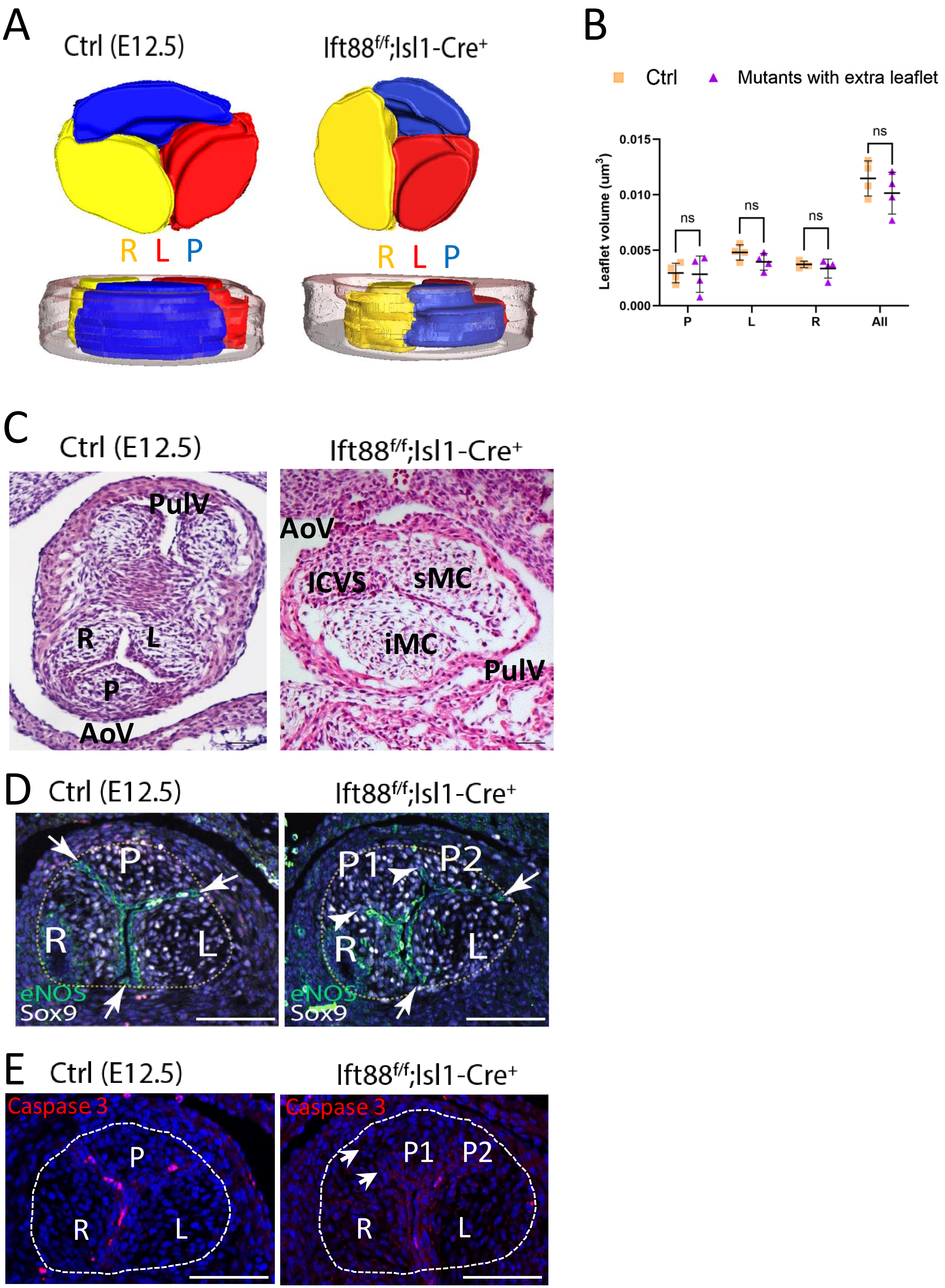

Supplementary Figure 7

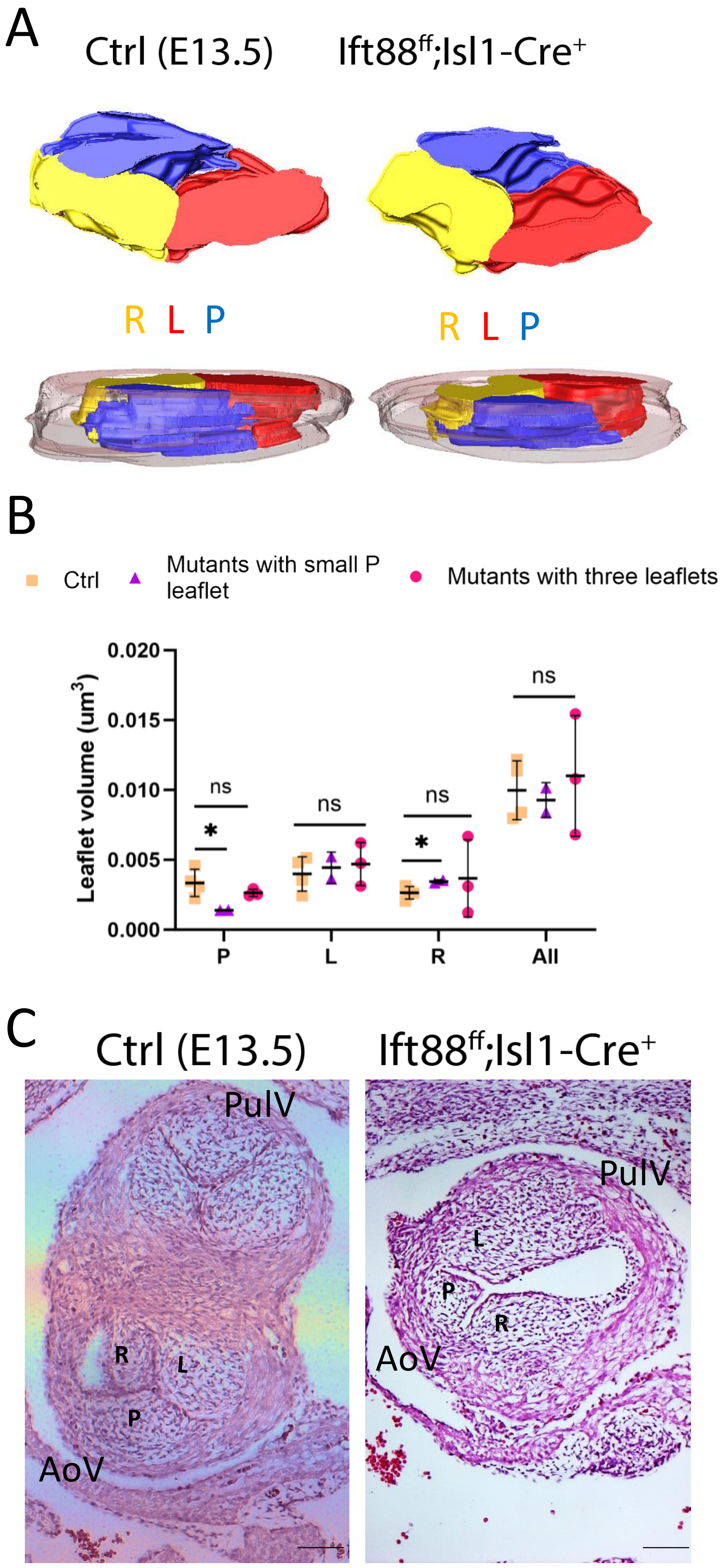

Supplementary Figure 8

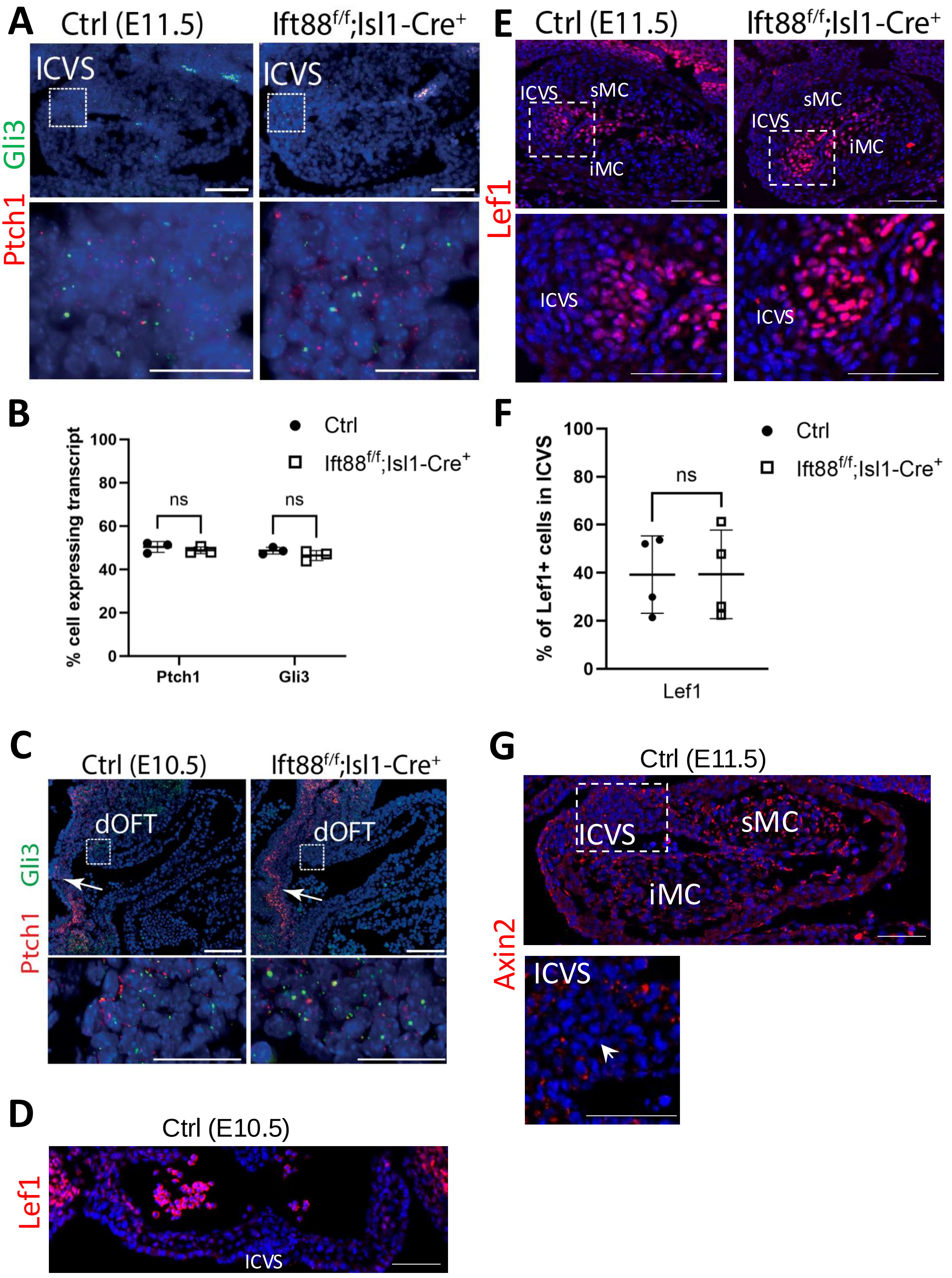

Supplementary Figure 9

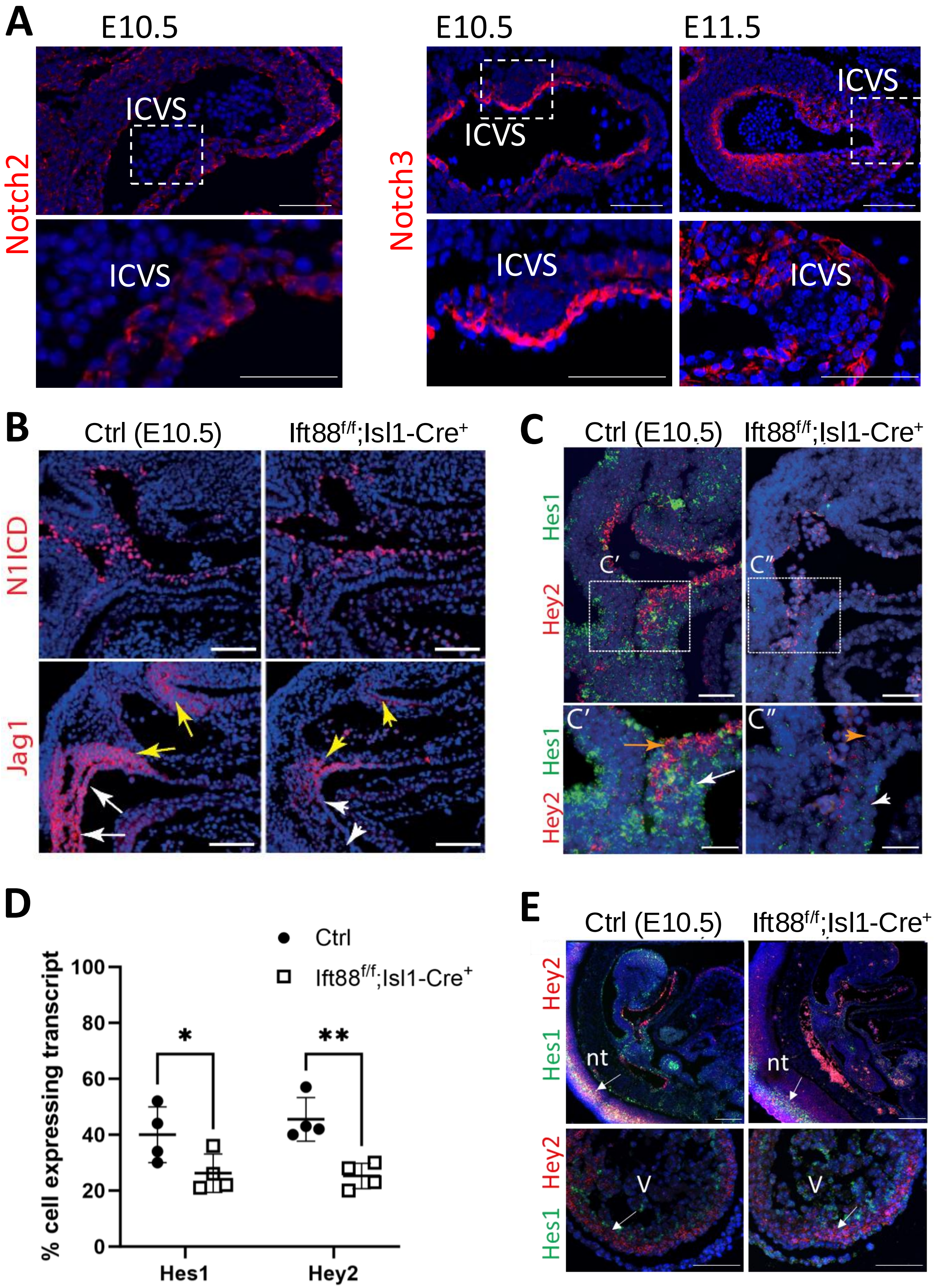

Supplementary Figure 10

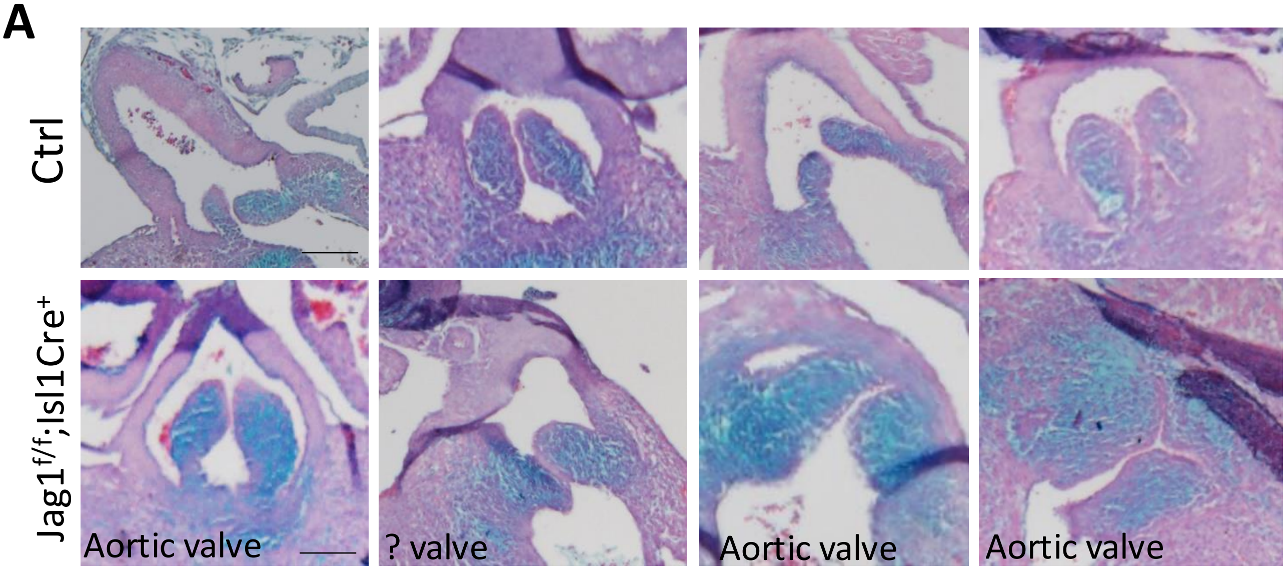
