## Supplementary figure legends for "Remodelling of supernumerary leaflet primordia leads to bicuspid aortic valve (BAV) caused by loss of primary cilia"

**Supplementary Figure 1:** **Primary cilia are found on the progenitor cells forming the arterial valves**. **A-A’)** Primary cilia labelled by acetylated-tubulin (green; arrows in A’) found on NCC (red) traced by Wnt1-Cre in the main outflow cushions at E10.5, n=3. **B-B’)** Primary cilia labelled by acetylated-tubulin (green; arrows in B’) found on EDC (red) traced by Tie2-Cre in the main outflow cushions at E10.5, n=3. **C)** Quantification of ciliated cells during the main stages of outflow cushion formation and remodelling showing the percentage of ciliated NCC and EDC. Most cells are ciliated at E10.5-E11.5 although this drops at E12.5 and is maintained at E13.5 to E15.5. These dynamic changes in cilia were similar between NCC and EDC, n=3 per genotype. Two-way ANOVA with Tukey test, Mean ± SD. ns: not significant, ****p<.0001. **D-D”)** Representative immunofluorescent images of primary cilia labelled by Arl13b (green; arrows in D’) found on undifferentiated SHF cells labelled by Isl1 (red) at E10.5. Cilia were absent from differentiated cardiomyocytes labelled by cTnI (white; arrowheads in D”), n=3. **E-E’)** Representative immunofluorescent images of primary cilia labelled by Arl13b (green) found on undifferentiated SHF cells labelled by Isl1 (red) at E11.5. Differentiated smooth muscle cells labelled by SM22a (white) did not carry a primary cilium (arrowheads in E’), n=3. In all cases, cell nuclei are labelled by DAPI (blue). **F-F’’)** Co-localisation of acetylated-tubulin and Ift88 in cilia in the OFT cushions, n=3.

NCC, neural crest cells; EDC, EndoMT derived cells.

Scale bar = 100μm

**Supplementary Figure 2:** **Additional defects observed in conditional Ift88 mutants**. **A)** Quantitative analysis of ciliary length and the percentage of ciliated cells in EDC from Ift88^f/f^;Tie2-Cre mutants and NCC from Ift88^f/f^;Wnt1-Cre compared to control littermates (Ctrl). This showed a significant reduction in both measurements in mutant embryos (n=3/group, Mean ± SD, two tail Unpaired *student t-*test, ***p0.001). **B)** Table showing the percentages of recovered Ift88^f/f^;Wnt1-Cre, Ift88^f/f^;Tie2-Cre and Ift88^f/f^;Tnnt2-Cre mutants at E15.5. Chi squared test showed no statistically significant differences between the expected and observed values**. C)** Representative immunofluorescent images of Ift88 (red) found on NCCs labelled by GFP (green) of the outflow tract in Wnt1-Cre^+^ embryos at E10.5, n=3. **D)** Analysis of the atrioventricular cushions and dorsal mesenchymal protrusion, as well as the outflow cushions, in Ift88^f/f^;Tie2-Cre^+^ mutants and control littermates at E11.5, using immunofluorescence to label the myocardium (cTnI; green) and mesenchymal cells (Sox9; red). This revealed that the atrioventricular cushions remained unfused in 2 of 6 mutant hearts examined, but no controls. In contrast, the dorsal mesenchymal protrusion was fused with the superior atrioventricular cushion in all cases and the outflow tract appeared normal in all hearts. **E)** Histological analysis of the pulmonary valve of Ift88^f/f^;TnnT2-Cre mutants at E15.5 suggested dysplasia of all three leaflets in one (1/8) mutant.

EDC, EndoMT derived cells; NCC, neural crest cells; AV, atrioventricular cushion; DMP, dorsal mesenchymal protrusion; OFT, outflow tract; L, left leaflet; R, right leaflet.

Scale bar= 100μm.

**Supplementary Figure 3:** **Limited abnormalities in Ift88^f/f^;Mef2c-AHF-Cre related to** **differences in Isl1-Cre and Mef2c-AHF-Cre expression patterns.** **A)** Histological analysis of Ift88^f/f^;Mef2c-AHF-Cre mutants and control littermates at E15.5 revealed no aortic valve malformations (n=8 for each genotype), but membranous VSD in 2/8 mutants (yellow arrow and arrowhead). **B)** Table showing the numbers of recovered mutants at E15.5 for Ift88^f/f^;Mef2c-AHF-Cre and Ift88^f/f^;Isl1-Cre crosses. Chi squared test showed no differences between expected and observed values. **C,D)** Spatiotemporal mapping of Mef2c-AHF-Cre and Isl1-Cre-derived cells in outflow tract development between E10.5 and E11.5 (n=3 per group and per stage). At E10.5, Isl1-Cre labels all cells in the ICVS (asterisk), and the endocardial cells found in the outflow region (green arrows), whereas there were some unlabelled cells in the ICVS (asterisk and white arrowheads) and the endocardial layer of the Mef2c-AHF-Cre outflow tract (short green arrows). At E11.5, Isl1-Cre continues to label all the ICVS cells (asterisk and white arrows) and the majority of the neighbouring endocardial cells (green arrows). In contrast Mef2c-AHF-Cre labelling excludes a subpopulation of cells in the ICVS (asterisk and short white arrows) and a subpopulation of the endocardial cells (short green arrows). **E)** Lineage tracing of Mef2c-AHF-Cre and Isl1-Cre at E12.5 (n=3 per group) continued to show differences in expression patterns. Although Mef2c-AHF-Cre and Isl1-Cre expression patterns were overlapping in all cells populating the left and right leaflets, there was a subpopulation of mesenchymal cells located in the distal part of the posterior leaflet that was labelled by Isl1-Cre, but not Mef2c-AHF-Cre (white arrows and arrowheads). The dashed boxed area shows the posterior leaflet at higher magnification. Endocardial cells overlaying the posterior leaflet were only labelled by Isl1-Cre (green arrows and arrowheads). **F)** The expression patterns of Isl1-Cre and Mef2c-AHF-Cre were similar in the anterior leaflet of the pulmonary valve at E15.5, n=3.

membVSD, membranous VSD; BAV, bicuspid aortic valve; ICVS, intercalated valve swellings; L, left leaflet; P, posterior; R, right leaflet; A, anterior.

Scale bar = 100μm

**Supplementary Figure 4:** **No evidence of changes in the extracellular matrix components of aortic valve in Ift88^f/f^;Isl1-Cre**^+^ **mutants. A)** Immunolabelling of periostin, collagen I and versican showed comparable expression patterns between controls and mutants at E18.5, n=3.

L, left leaflet; P, posterior; R, right leaflet.

Scale bar =100μm

**Supplementary Figure 5:** **Neither cardiomyocytes nor SMC differentiate prematurely in Ift88^f/f^;Isl1-Cre**^+^ **mutants.** **A)** Immunolabelling of cardiomyocytes (labelled by cardiac troponin I; cTnI; white) and SHF (labelled by Isl1; red) showed comparable patterns in the distal outflow tract (arrows) in Ift88^f/f^;Isl1-Cre mutants and controls at E10.5 (n=3 per genotype), suggesting that the time course of differentiation of SHF cells to cardiomyocytes was similar in both cases. **B)** Immunolabelling for SMC (labelled by SM22α; white; arrowheads) with the ICVS (bracket) labelled by Isl1 (red, arrows) showed no differences in the pattern of smooth muscle cells between Ift88^f/f^;Isl1-Cre^+^ mutants and controls at E11.5, suggesting that the time course of differentiation of SHF cells to SMC was similar in both cases (n=3 for each genotype). **C)** 3D reconstructions, produced from anti-eNOS immunofluorescence images, revealed the extent of the endocardium is foreshortened in the mutants (arrowheads) compared to controls (arrows) with the supernumerary ICVS (bracket) is juxtaposition with the adjacent cushion. **D)** Immunolabelling for cell death using Caspase-3 (red) revealed no evidence of cell death in the region of mesenchymal continuity (arrowheads) between the ICVS and adjacent main cushion in the mutants.

A-ICVS, anterior (pulmonary) intercalated valve swelling; P-ICVS, posterior (aortic) intercalated valve swelling; iMC, inferior main cushion; sMC, superior main cushion; Ctrl, control; OFT, outflow tract.

Scale bar =100μm

**Supplementary Figure 6:** **Ift88^f/f^;Isl1-Cre mutant with three aortic valve leaflets at E12.5 have subtle defects in the shape of the leaflets.** **A)** 3D reconstructions of Ift88^f/f^;Isl1-Cre mutants with three aortic valve leaflet primordia at E12.5 showed slight changes in the size of the posterior leaflet compared to controls, n=2/6. **B)** Quantification of the leaflet volume in Ift88^f/f^;Isl1-Cre^+^ mutants at E12.5 were comparable to controls, n=4. Two-way ANOVA with Šídák test, Mean ± SD. ns: not significant. **C)** Delay in the development of valve leaflets and septation of the outflow tract was observed in 1/6 Ift88^f/f^;Isl1-Cre^+^ mutants collected at E12.5. **D)** Immunolabelling of cushion mesenchyme (labelled by Sox9; white) and the endocardium overlying each leaflet primordium (labelled by eNOS; green) showed that all three commissures were close to the arterial wall in controls (arrows) whereas at least one commissure was displaced towards the lumen in Ift88^f/f^;Isl1-Cre^+^ mutants (arrowheads; n=3/3. **E)** Representative immunofluorescent images of caspase-3 (red) showed no evidence of cell death in the region of fusion seam (arrowheads) between the aortic valve leaflets in the mutants, n=3.

AoV, aortic valve; PulV, pulmonary valve; P, posterior leaflet; leaflet; R, right leaflet; L, left leaflet; iMC, inferior main cushion; sMC, superior main cushion.

Scale bar =100μm

**Supplementary Figure 7: Posterior leaflet appeared smaller even in those Ift88^f/f^;Isl1-Cre mutants that seemed apparently normal at E13.5.** **A)** 3D reconstructions of the aortic valve leaflet primordia at E13.5 in Ift88^f/f^;Isl1-Cre^+^ mutants revealed that the posterior leaflet appeared slightly smaller compared to controls even in mutants that looked apparently normal from histological sections, n=3/7. **B)** Quantification of the leaflet volume in Ift88^f/f^;Isl1-Cre^+^ mutants at E13.5 were comparable to controls for the group of mutants with three leaflets (n=3), but significantly different between posterior and right leaflets for the group of mutants with small posterior leaflet (n=2). Two-way ANOVA with Šídák test, Mean ± SD. ns: not significant, *p<.05**. C)** Delay in the septation of the outflow tract was observed in 2/7 Ift88^f/f^;Isl1-Cre^+^ mutants collected at E13.5.

AoV, aortic valve; PulV, pulmonary valve; L, left leaflet; R, right leaflet; P, posterior leaflet.

Scale bar =100μm

**Supplementary Figure 8: Unaltered Hedgehog or Wnt signalling pathways in the ICVS of Ift88^f/f;^Isl1-Cre mutants. A)** Representative RNAscope images (n=3 per genotype) for Ptch1 and Gli3 expression in the ICVS at E11.5 showed no reproducible differences between Ift88^f/f^;Isl1-Cre^+^ mutants and controls. **B)** Quantification of Ptch1 and Gli3 expression in the ICVS indicated no significant differences in the number of positive cells expressing each marker between mutants and control littermates. Two-way ANOVA with Šídák test, Mean ± SD, ns: not significant. **C)** Representative RNAscope images (n=4 per genotype) E10.5 revealed that although Ptch1 and Gli3 expression levels in the outflow tract were comparable between Ift88^f/f^;Isl1-Cre^+^ mutants and controls, Ptch1 was upregulated in the pharyngeal and paraxial mesoderm (arrow) of the mutants compared to controls. **D)** Immunolabelling of the Wnt signalling marker Lef1 (red) at E10.5 showed no expression of this marker in the ICVS at this stage. **E)** Representative immunofluorescent images of Lef1 at E11.5 showed comparable expression level between Ift88^f/f^;Isl1-Cre^+^ mutants and controls, n=4. **F)** Quantification of Lef1 expression in the ICVS indicated no significant differences in the percentage of positive cells expressing this marker between mutants and control littermates. Unpaired *t-test*, Mean ± SD, ns: not significant. **G)** Immunolabelling of the Wnt signalling marker, Axin2 (red), at E11.5 showed no expression of this marker (arrowhead) in the ICVS.

ICVS, intercalated valve swellings; iMC, inferior main cushion; sMC, superior main cushion.

Scale bar =100μm

**Supplementary Figure 9: Disruption of Notch signalling in the dorsal pericardial wall and distal outflow tract walls of the Ift88^f/f;^Isl1-Cre mutants.**

**A)** Immunolabelling of Notch2 at E10.5 and Notch3 at both E10.5 and E11.5 revealed no expression of these markers in the ICVS, n=3 per marker per stage. **B)** Immunofluorescence analysis of N1ICD expression at E10.5 showed comparable patterns between all four Ift88^f/f^;Isl1-Cre mutants and their littermate controls. Immunofluorescence for Jag1 at E10.5 showed a marked reduction in the expression level of Jag1 in distal outflow tract (yellow arrowheads) and dorsal pericardial wall (white arrowheads) in 4/4 Ift88^f/f^;Isl1-Cre mutants compared to littermate controls. **C)** Representative RNAscope images (n = 4/group) of E10.5 control and Ift88^f/f^;Isl1-Cre mutants showed reduction in Hes1 expression in the dorsal pericardial wall of mutants (green, white arrowhead in G’’), and diminished Hey2 expression in the outflow tract endocardium of mutants (red, orange arrowhead in G’’) of 3/4 Ift88^f/f^;Isl1-Cre mutants. **D)** Quantification of the number of cells expressing Hes1 and Hey2 in the distal outflow tract identified a significant reduction in the number of positive cells in mutants compared to controls at E10.5. Two-way ANOVA with Šídák test, Mean ± SD. *p=.047, **p=.005. **E)** Representative images showing the reduction in the expression level of Hey2 (red) and Hes1 (green) in Ift88^f/f^;Isl1-Cre mutants was restricted to the SHF and outflow tract, as the expression pattern of Hes1 in the neural tube (nt, arrow) and Hey2 in ventricle (V, arrow) were comparable between the mutants and controls, n=4.

ICVS, intercalated valve swellings; nt, neural tube; V, ventricle.

Scale bar =100μm

**Supplementary Figure 10: Abnormal arterial valves in Jag1^f/f^;Isl1-Cre mutants. A)** Representative images of the range of arterial valve abnormalities found in the Jag1^f/f^;Isl1-Cre mutants (n=4) compared to littermate controls at E14.5.

Scale bar =100μm
